# Wastewater and colloidal extracts of wastewater-irrigated soils select for resistant *Acinetobacter baylyi* beyond what measured antibiotic concentrations predict

**DOI:** 10.64898/2026.05.12.724625

**Authors:** Katharina Axtmann, Christina Paffenholz, Alina Auerhammer, Ana Karen Michel-Farias, Benjamin Justus Heyde, Lena Marie Coppers, Melanie Braun, Arne Kappenberg, Ines Mulder, Susanne Brüggen, Christina Siebe, Wulf Amelung, Jan Siemens, Gabriele Bierbaum

**Affiliations:** Institute of Medical Microbiology, Immunology and Parasitology, University Hospital Bonn, Venusberg-Campus 1, 53127 Bonn, Germany; Ruhr University Bochum, Institute of Geography, Unit Soil Sciences and Soil Resources, Universitätsstraße 150, 44801 Bochum, Germany; Institute of Crop Science and Resource Conservation, Soil Protection and Ecosystem Health, University of Bonn, Nussallee 13, 53115 Bonn, Germany; North Rhine-Westphalia Office of Nature, Environment and Climate, Leibnizstraße 10, 45659 Recklinghausen, Germany; Instituto de Geología, Universidad Nacional Autónoma de México, Mexico City, Mexico; Institute of Soil Science and Soil Conservation, iFZ Research Center for BioSystems, Land Use and Nutrition, Justus Liebig University Giessen, Heinrich-Buff-Ring 26-32, 35392 Giessen, Germany

**Keywords:** wastewater irrigation, antibiotic resistance selection, soil colloids, pharmaceutical residues, minimal selective concentrations

## Abstract

Numerous studies have shown that the abundance of antibiotic-resistant bacteria (ARBs) or antibiotic-resistance genes (ARGs) in soil increases after irrigation with wastewater. However, it is unclear whether this increase is due to the selection effects of pharmaceutical residues in the irrigation water or the continuous introduction of ARBs and ARGs with the wastewater. Further, it is unclear how the binding of antibiotics to natural colloids (1-1000 nm) affects their biological effects compared to truly dissolved substances (< 1 nm). We conducted competition experiments with resistant and susceptible *Acinetobacter baylyi* BD413 strains in wastewater, as well as in colloidal and truly dissolved extracts of soils irrigated with wastewater. Although the concentrations of our six target antibiotics were far below the measured minimum selective concentrations of the tested strains, we demonstrate that the resistant strain was favored in the wastewater and the colloidal extracts. In contrast, the truly dissolved fractions exhibited weaker and more variable selective effects. A non-targeted analysis revealed the presence of 82 additional substances in our extracts, including further antibiotics, pesticides, and different non-antibiotic drugs that may influence the selection of our resistant *A. baylyi* BD413 strain. Our findings suggest that antibiotic resistance is selected for in wastewater and wastewater-irrigated soils. This cannot be explained by antibiotic concentrations alone, but may also arise from the effects of complex mixtures of co-occurring contaminants, particularly those associated with colloidal particles.

## 1. Introduction

Antibiotic therapy is one of the most transformative advances in modern medicine, significantly improving human survival and quality of life. However, since the discovery of penicillin in 1928 [1] and its widespread clinical use in the 1940s, the emergence and spread of antibiotic-resistant bacteria (ARB) have become a major public health concern [2,3]. Recent global estimates suggest that antibiotic resistant bacteria (ARBs) were associated with approximately 4.71 million deaths in 2021, including 1.14 million deaths directly attributable to antimicrobial resistance [4]. Projections indicate an ongoing increase, with ARB-related mortality expected to reach 8.22 million cases per year by 2050, 1.91 million of which will be directly attributable to ARB infections [4]. These trends underscore the significant threat that ARBs pose to the sustainability of healthcare systems and the efficacy of modern therapeutic interventions.

The selection and spread of ARBs that cause disease in humans or domestic animals is primarily driven by the administration of antibiotics in clinics or in livestock breeding. However, the role of the environment as a reservoir of ARBs and antibiotic resistance genes (ARG) has received increasing attention in recent years in the context of the One Health concept [5,6]. Since most antibiotics are not fully metabolized, residues are excreted into wastewater through urine and feces from humans and animals [7–9]. Improper disposal of antimicrobials [10] and effluents from production facilities also contribute to the presence of antibiotic residues in wastewater and the environment [11,12].

In wastewater systems, and particularly within wastewater treatment plants (WWTPs), the horizontal transfer of ARGs among bacteria occurs at elevated rates due to the simultaneous presence of various antibiotics and ARBs [13]. Globally, the primary design of WWTPs focuses on removing solids, nutrients (nitrogen and phosphorus), and dissolved matter since the effluents are afterwards discharged into surface water bodies. However, they are considerably limited in their ability to eliminate organic micropollutants, such as pharmaceuticals. Consequently, pharmaceutical residues, including antibiotics, are discharged into the environment with WWTP effluents [14,15].

In many regions of the world, primarily arid and semiarid regions, both treated and untreated wastewater is used for crop irrigation to prevent water shortages and recycle nutrients [16]. This poses potential risks to field workers and consumers of the produce [17–19]. It is expected that around one million hectares globally will be irrigated with treated wastewater, while approximately 20 million hectares are currently irrigated with untreated, partially treated, or diluted wastewater [20]. Due to the expected changes in precipitation and increase in droughts resulting from climate change, the demand to irrigate agricultural fields will most likely increase globally [21–24].

Long-term irrigation with wastewater can result in the accumulation of pharmaceuticals in soils. Studies have shown that concentrations of antibiotics, such as sulfamethoxazole (SMX), trimethoprim (TRM), and ciprofloxacin (CIP), increase with prolonged irrigation [25–27]. These antibiotics can be taken up by crops and consumed by humans or animals [28–30], thus transporting antibiotics by irrigation to the soil, then to plants and back to humans. Additionally, the accumulation of antibiotic residues may lead to increased selection pressure, resulting in elevated abundances of ARBs and ARGs [25,27,31]. Typically, the accumulated concentrations do not reach the minimum inhibitory concentrations (MICs) of susceptible bacteria [32,33]. However, they can reach minimal selective concentrations (MSCs) and therefore, confer growth advantages to ARBs [34]. Combinations of different antibiotics and other pharmaceuticals (e.g., disinfectants or heavy metals) can further lower MSCs, co-select for ARBs, and increase horizontal gene transfer (HGT) [35–38].

The Mezquital Valley in Hidalgo, Mexico, is one of the largest regions in the world that uses wastewater for irrigation. Since irrigation with untreated wastewater from the metropolitan areas of Mexico City (mostly by flooding) commenced more than 100 years ago, the irrigated area has increased to more than 90,000 hectares, allowing crop cultivation in this semi-arid area with low rainfall [39,40]. In 2017, the Atotonilco WWTP began operating and treats around 60% of the wastewater from the Mexico City metropolitan areas. The remaining wastewater is used without prior treatment for irrigation in the Mezquital Valley or enters the Tula River [41]. Previous studies have documented the extent of contamination in the Mezquital Valley’s irrigation wastewater, as well as the accumulation of pollutants in soil, groundwater, and crops [25,42–47] inducing an increase in ARBs and ARGs [25,31,48].

To reach the Sustainable Development Goal (SDG) 6, “Clean Water and Sanitation” [49], more investments in wastewater collection and treatment will be made. This will induce a shift from using untreated wastewater for irrigation to using treated wastewater [50]. While many studies have compared the risks of wastewater irrigation with rainfed agriculture or groundwater irrigation, only a very limited number of studies investigated the consequences of transitioning from irrigating with untreated wastewater to treated wastewater [16]. Soil organic matter (SOM) is a major sorbent of pollutants. A decline in SOM content due to the shift to treated wastewater irrigation may, thus, release contaminants in the curse of SOM degradation and reduce the soil’s ability to retain contaminants. This could enhance the mobilization of legacy pollutants, which could leach into groundwater, be taken up by crops, or affect ARBs or ARGs selectively [16]. Carillo et al. (2016) [51] already demonstrated the desorption of SMX following simulated irrigation with treated wastewater in Leptosols and Phaeozems of the Mezquital Valley. In clay-rich Vertisols, this effect was not observed, because similar to SOM, clay is a major sorbent for SOM. Heyde et al. (2025) [52] demonstrated the solubilization of toxic metals Cr and Ni from Mezquital Valley soils with a long-term history of irrigation with untreated wastewater upon irrigation with treated wastewater.

Clay and SOM may be present in soil in the form of soil colloids, i.e., as particles in the range of 1 to 1000 nm [53]. Natural colloids occurring in rivers, soil, but also in wastewater consist mainly of secondary minerals from weathering, of organic material, and of organo-mineral associations [54–57]. During wastewater treatment, a fraction of these colloids are removed; nevertheless, some colloids escape WWTPs and are hence frequently found in wastewater effluents (e.g., reviewed in Shon et al. (2006) [58]). In soil, the chemical, mineralogical, and organic composition of colloids is highly dependent on soil properties and environmental conditions. Colloids are important binding partners for pharmaceuticals in wastewater and also later in the environment, i.e., rivers and soil, after discharge from WWTPs. Higher partitioning coefficients for pharmaceuticals to colloids as compared to dissolved organic carbon, total suspended organic matter or the bulk sediments have been reported (reviewed in Bagnis et al. (2018) [59]). Preferred binding of pharmaceuticals is caused by the small size of the particles, resulting in high specific surface areas of > 10 m^2^/g, high surface charge, and high reactivity of colloids [60,61]. Hence, depending on their physicochemical properties, large fractions of pharmaceuticals are bound to colloids in wastewater effluent and rivers. For example, Maskaoui & Zhou [62] found that 7 to 84% of pharmaceuticals (propranolol, SMX, carbamazepine, indomethacin and diclofenac) were associated with colloids in a study of UK rivers and groundwater as well as wastewater effluents, suggesting that these colloids play a major role for the fate and transport of those compounds in WWTPs.

Therefore, our overall aim was to study the effect of colloidal binding on the effectiveness on antibiotics in wastewater and wastewater-irrigated soils. In detail, we formulate the following hypotheses: (i) the total amount of selecting pharmaceuticals in wastewater and soils can trigger the selection of ARBs, even though (ii) the binding of antibiotics to soil colloids will reduce their effectiveness and hence will cause a lower selection of ARBs than antibiotics in the truly dissolved phase. Furthermore, we hypothesize that (iii) a shift from irrigation with untreated wastewater to treated wastewater may cause a stronger selection of ARBs by solubilizing accumulated antibiotics from the soil [16]. To test these hypotheses, we performed competition experiments using fluorescently marked *Acinetobacter baylyi* BD413 strains (resistant and susceptible, see Schuster et al. (2022) [63]) in wastewater of different qualities, and in colloidal and truly dissolved extracts from three different soils from the Mezquital Valley, irrigated with WWTP influent and effluent.

## 2. Material & Methods

### 2.1 Minimal inhibitory concentrations and minimal selective concentrations

MICs for the individual strains and MSCs for *A. baylyi* BD413 mCherry 652 or *A. baylyi* BD413 mCherry 777 each in competition with *A. baylyi* BD413 GFP were determined for SMX, azithromycin (AZI), TRM and erythromycin (ERY) as described in Schuster et al. (2022) [63].

### 2.2 Soil & wastewater samples-incubation experiment

Soil samples were taken from three different soil reference groups in the Mezquital Valley in Mexico that had been irrigated with untreated wastewater for more than 60 years, classified according to WRB as Vertisol, Phaeozem, Leptosol [64] (in the following referred to as soil type). Four different fields were sampled per soil type. Eight samples were collected from four locations within each field, which were then mixed to create a composite sample. Composite wastewater samples were taken from the influent and effluent of the Atotonilco WWTP [65]. All soil and water samples were stored and transported under cooled conditions (< 8 °C) until the start of the experiment.

Before beginning the incubation experiment, different chemical properties and the concentrations of the following antibiotics in the wastewater samples were determined: CIP, SMX, TRM, AZI, clindamycin (CLI), anhydroerythromycin (A-ERY), and ERY. These antibiotics were selected due to their longevity and previous consumption data, furthermore, a broad spectrum of chemical classes was to be covered [65]. The concentrations of the following quaternary alkyl ammonium compounds (QAACs) were also determined: alkyltrimethyl ammonium compound (ATMAC)-C8, ATMAC-C10, ATMAC-C12, ATMAC-C14, ATMAC-C16, benzylalkyldimethyl ammonium compound (BAC)-C8, BAC-C10, BAC-C12, BAC-C14, BAC-C16, BAC-C18, dialkyldimethyl ammonium compound (DADMAC)-C8, DADMAC-C10, DADMAC-C12, DADMAC-C14, DADMAC-C16, and DADMAC-C18 (see Table S1 of Soufi et al. (2025) [65]). Various soil properties were also analyzed (see Table S2 of Soufi et al. (2025) [65]). The sampling process and the chemical analysis of the samples are described in more detail in Soufi et al. (2025) [65] and Heyde et al. (2025) [52].

Each composite soil sample was divided into four subsamples, which were then mixed with one of the following: (I) unspiked WWTP influent, (II) unspiked WWTP effluent, (III) WWTP influent spiked with the following antibiotics and disinfectants: A-ERY, AZI, CIP, CLI, ERY, SMX, TRM, ATMACs, DADMACs and BACs), or WWTP effluent spiked with the same antibiotics and disinfectants. The spiking was performed to establish the 500-fold concentrations of the pharmaceuticals in comparison to the concentrations measured in the WWTP influent. Spiking was performed to simulate repeated irrigation scenarios and enable tracking of the dissipation of wastewater-derived compounds over time at levels exceeding the limits of quantification.

The water content of the soil samples was adjusted to 80% of their water-holding capacity using distilled water. Subsequently, 200 g of each soil (dry weight equivalent) was mixed with 25 mL of the respective wastewater type. Afterwards, the soil microcosms were covered with sterile surgical masks to minimize contamination from dust or insects. The microcosms were then incubated for up to four weeks in a climatic chamber maintained at 95% relative humidity and 22 °C. Samples were withdrawn after four days and four weeks.

### 2.3 Colloidal and truly dissolved extracts and antibiotic concentrations

According to Braun et al. (2025) [66], samples were frozen after the experiment and thawed before analysis. Extraction and fractionation of the colloidal and truly dissolved phase was conducted as follows: In a first step, so-called water-dispersible colloids were extracted from soil by shaking 10 g of soil with 60 mL of Millipore water for 24 h. Afterwards, to remove soil particles > 1,000 nm, extracts were centrifuged at 1,800 rpm for 5 min. and the supernatant was transferred to ultracentrifuge vials for further separation of the truly dissolved phase. The formed pallet was discarded. The removed and transferred supernatant in the ultracentrifuge tubes was filled with Millipore water to the maximum filling volume of 66 mL. The samples were then centrifuged at 45,000 rpm for 26 h. The resulting supernatant after ultracentrifugation represents the truly dissolved fraction and the remaining pellet the colloid fraction < 450 nm.

Subsequently, the truly dissolved fraction (supernatant) was transferred and completely evaporated. The sample was then re-dissolved in methanol and passed through a filter (0.22 µm). A further complete evaporation and re-dissolution of the sample was carried out with 0.225 µL H_2_O/ MeOH (90:10; v:v) and 0.025 µL internal standard solution in GC vials. The antibiotics in the truly dissolved fraction from these suspensions were measured on an Acquity H-class UPLC (Waters, Eschborn, Germany) according to a modified method of Dalkmann et al. (2012) [25] and Kappenberg & Juraschek [67]. The colloid fraction (remaining pellet after ultracentrifugation) was further extracted in order to isolate the antibiotics associated with the colloid fraction. Therefore, the pellet was mixed with 25 mL of the extraction solvent MeOH/ H_2_O/ ACN/ Acetic Acid (0.01%), (50:25:24:1; v:v:v:v) and placed on the horizontal shaker for 30 min. The sample was then placed in an ultrasonic bath for 10 min and afterwards centrifuged at 4,000 rpm for 10 min. The supernatant was carefully removed and the centrifuge tube was shaken again with 20 mL of extraction solvent and centrifuged as described above. The second supernatant was removed and mixed with the first supernatant. The entire supernatant was completely evaporated and re-dissolved with 1 mL H_2_O/ MeOH (90:10, v:v). The sample was then centrifuged again at 10,000 rpm for 10 min to remove potential contamination. For the measurement, 200 µL of the final sample was transferred and measured by LC-MS/MS.

### 2.4 Competition experiments in wastewater and soil extracts

The soil extracts (see 2.2) were used to perform competition experiments with antibiotic-resistant and susceptible *A. baylyi* BD413 strains labeled with either GFP or mCherry (for a detailed description see Schuster et al. (2022) [63]). For the competition experiments, 4.5 mL of each extract was mixed with 0.5 mL of Müller-Hinton broth (MHB) in order to provide nutrients for growth and then inoculated with a 1:1 mixture (5×10^5^ cells/mL) of resistant and susceptible *A. baylyi* BD413 cells (with different fluorescence). Subsequently, the extracts were incubated overnight at 37 °C and 170 rpm. Afterwards, the extracts were diluted in a 1:10^−5^ ratio with sterile 0.9% NaCl and plated on MH agar containing 150 µg/mL spectinomycin. After incubating for 48 h at 30 °C, the colony forming units (CFUs) of the respective resistant and susceptible *A. baylyi* BD413 strains were counted to calculate their ratio. Equally structured competition experiments were performed with the strains in the four different wastewater types.

Furthermore, *A. baylyi* BD413 GFP pHHV216 and *A. baylyi* BD413 mCherry pHHV216 were used for the competition experiments in extracts that had shown a strong selection for *A. baylyi* BD413 mCherry 652. Both strains harbor the pHHV216 resistance plasmid [68], which confers resistance to sulfonamides, (*sul2*), tetracycline (*tetH*), chloramphenicol (*florR*), and streptomycin (*strA* and *strB*). However, due to fitness costs they also have a decreased growth rate compared to their parent strains and also compared to *A. baylyi* BD413 mCherry 652 [63]. As a control, the competition experiments with all strain combinations were also performed in 1:10-diluted MHB without any antibiotics. At least three biological replicates were conducted for all experiments.

### 2.5 Statistical analysis

Statistical analyses were conducted in R (version 4.4.1) using the packages tidyr, dplyr, *car*, *broom*, and *readODS*. Data were imported from an OpenDocument Spreadsheet and converted into long format with the percentage of *A. baylyi* BD413 mCherry 652 CFUs from the total CFUs as the response variable. The categorical factors included soil type, wastewater type, spiking, and sampling date. For each factor, a one-way ANOVA was performed to test for effects on mCherry fluorescence. Model assumptions were evaluated using the Shapiro–Wilk test for normality and Levene’s test for homogeneity of variances. Factors with *p* < 0.05 were considered statistically significant.

### 2.6 Rebuilt antibiotic concentrations

To determine whether the observed selection effects in the wastewater or the soil extracts were caused by the set of antibiotics, the measured concentrations (see 2.2) of randomly selected extracts were reconstituted into 1:10-diluted MHB. The concentrations of the six target antibiotics measured in the WWTP influent and effluent as well as the concentrations that were spiked into the wastewater (Table 1) were also reconstituted. Afterwards, competition experiments with *A. baylyi* BD413 mCherry 652 in combination with *A. baylyi* BD413 GFP were performed as described above. As a control, the competition experiment was performed in 1:10-diluted MHB without any antibiotics. At least three biological replicates were conducted for all experiments.

**Table 1:** Measured and spiked concentrations [µg/L] of the six target antibiotics from the WWTP influent and effluent, that were reconstituted in 1:10-diluted Müller-Hinton broth. *: values determined and spiked by Soufi et al. (2025) [65]; **: as CLI was not detected, the spiked concentration was calculated on the base of concentrations published by Siemens et al. (2008) [47]; QL: quantification limit; DL: detection limit.

| Antibiotic [$\mu\text{g/L}$] | WWTP influent* | WWTP effluent* | Spiked concentrations* |
| --- | --- | --- | --- |
| SMX | 3.19-3.99 | 2.82-3.78 | 1,995 |
| TRM | 1.1-1.3 | 1.2-1.4 | 650 |
| ERY | 0.21 (< QL) -0.37 | 0.17-0.25 (< QL) | 185 |
| CLI | < DL | < DL | 60** |
| CIP | 2.35-3.39 | 1.94-2.27 | 1,695 |
| AZI | 0-1.39 | 0-1.33 | 695 |

### 2.7 Non-targeted analysis

Six selected colloidal extracts (three strongly selecting and three weakly selecting, all unspiked) were examined in a non-targeted analysis to investigate further potential selecting substances apart from the six target antibiotics. The analysis was performed as described by Brüggen et al. (2018) [69]. A suspect screening approach was performed to get qualitative information about the appearance of further micropollutants in the samples. Therefore, no reference standard was needed, and a database of currently circa 3,500 compounds was scanned.

## 3. Results

### 3.1 Minimal inhibitory concentrations and minimal selective concentrations

The determined MIC and MSC values for *A. baylyi* BD413 mCherry 652 and *A. baylyi* BD413 mCherry 777 are listed in Table 2 and 3. For the determination of the MSCs, competition experiments with the susceptible *A. baylyi* BD413 GFP and one resistant strain (*A. baylyi* BD413 mCherry 652 or *A. baylyi* BD413 mCherry 777) were performed in MHB in the presence of different antibiotic concentrations as described in Schuster et al. (2022) [63].

**Table 2:** MIC values [µg/L] of the tested *A. baylyi* BD413 strains. *: values determined by Schuster et al. (2022) [63].

| *A. baylyi* strain | TRM | ERY | AZI | SMX* | CIP* | CLI* |
| --- | --- | --- | --- | --- | --- | --- |
| | $\mu\text{g/L}$ | | | | | |
| **GFP** | 32,000 | 2,000 | 250 | 3,000* | 20* | 8,000* |
| **mCherry 652** | > 64,000 | 16,000 | 1,000 | 16,000* | 313* | 64,000* |
| **mCherry 777** | > 64,000 | 8,000 | 250 | 128,000* | 313* | 64,000* |

**Table 3:** MSC values [µg/L] of the tested *A. baylyi* BD413 strains. For the MSC determination, *A. baylyi* BD413 mCherry 652 and *A. baylyi* BD413 mCherry 777 were both grown in competition with *A. baylyi* BD413 GFP. *: values determined by Schuster et al. (2022) [63]. **: MIC of *A. baylyi* BD413 GFP.

| Antibiotic | Resistant strain | MSC [$\mu\text{g/L}$] | MSC/MIC** |
| --- | --- | --- | --- |
| TRM | mCherry 652 | 1799.7 | 0.06 |
|  | mCherry 777 | 2845.2 | 0.08 |
| ERY | mCherry 652 | 255.5 | 0.13 |
|  | mCherry 777 | 559.7 | 0.07 |
| AZI | mCherry 652 | 36.7 | 0.15 |
| SMX | mCherry 652 | 341.6 | 0.02 |
| CIP* | mCherry 652 | 16.7* | 0.45* |
|  | mCherry 777 | 51.5* | 1.39* |
| CLI* | mCherry 652 | 687.8* | 0.09* |
|  | mCherry 777 | 1000.2* | 0.13* |

### 3.2 Wastewater favors growth of the resistant *A. baylyi* BD413 mutant

In a first step, competition experiments were performed in the spiked and unspiked influent and effluent of the WWTP with *A. baylyi* BD413 mCherry 652, *A. baylyi* BD413 mCherry 777 or fluorescently labelled *A. baylyi* BD413 strains carrying the pHHV216 resistance plasmid. The two mutants (mCherry 652) and (mCherry 777) have a reduced susceptibility to several antibiotics due to mutations in (mCherry 777) or a deletion of (mCherry 652) the AdeN regulator of the AdeIJK efflux pump. *A. baylyi* BD413 mCherry 777 also carries mutations in a further two-component system (BmfS). All resistant strains were grown in competition with a susceptible *A. baylyi* BD413 strain with the opposite fluorescent marker, starting with a 1:1 ratio of both strains at the beginning of the incubation. As a control, the competition experiments were also performed in 1:10-diluted MHB without added antibiotics.

After the incubation and plating, both resistant mutant strains of *A. baylyi* BD413 clearly outcompeted the susceptible *A. baylyi* BD413 GFP strain, when grown in the spiked wastewater samples, reaching up to 80% of the total CFUs (wastewater, Figure 1). While *A. baylyi* BD413 mCherry 652 also outcompeted the susceptible strain in the unspiked samples, *A. baylyi* BD413 mCherry 777 only reached 8% or 19%, respectively, when grown in the unspiked influent or effluent. The strains of *A. baylyi* BD413 carrying the pHHV216 resistance plasmid also did not outcompete the susceptible strain. In the controls in media without antibiotics, the resistant strains reached 40%, 4%, or 15%, respectively, for *A. baylyi* BD413 mCherry 652, *A. baylyi* BD413 mCherry 777, or the pHHV216 plasmid carrying strains.

**Figure 1:**
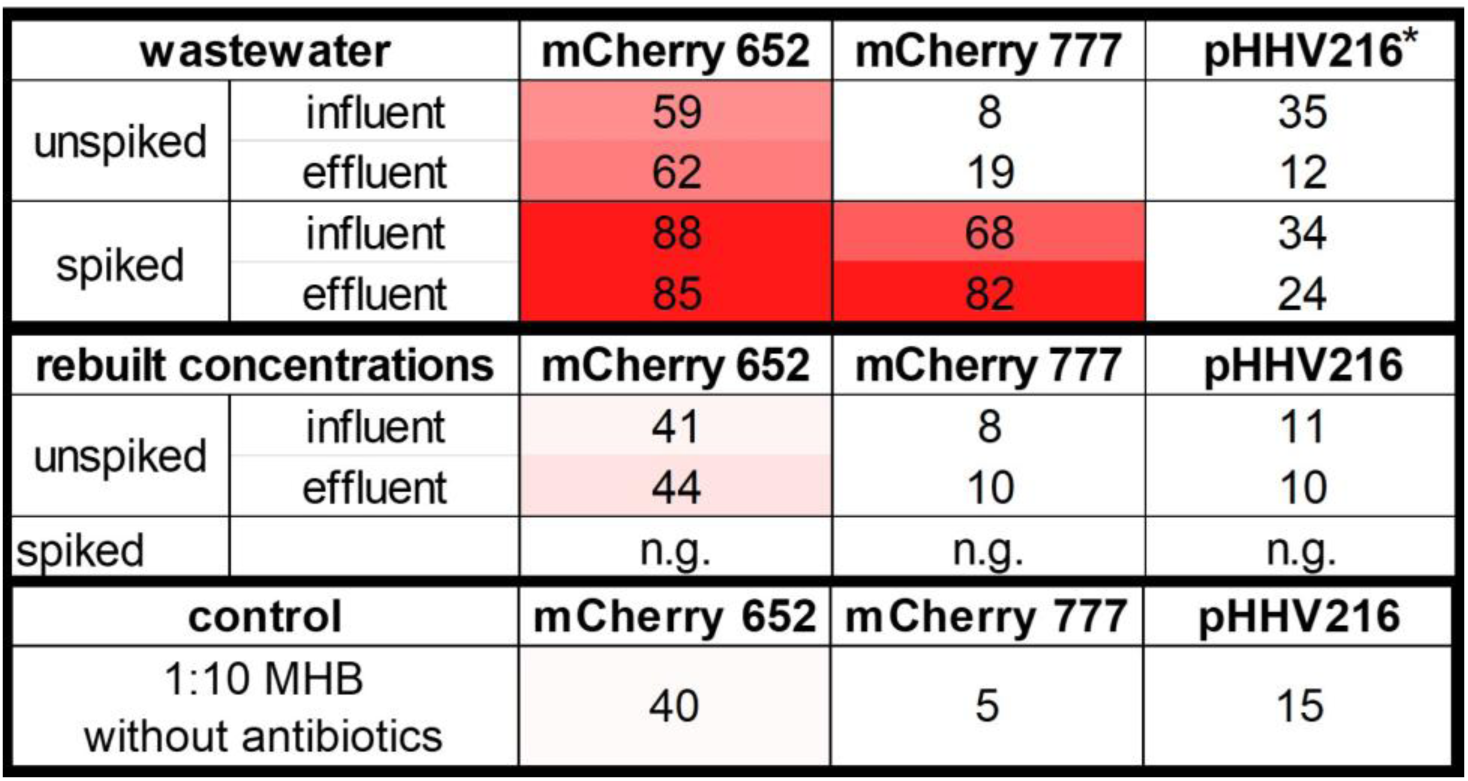
Heatmap showing the percentage of growth (CFUs) of *A. baylyi* BD413 mCherry 652, *A. baylyi* BD413 mCherry 777 or fluorescently labelled *A. baylyi* BD413 strains carrying the pHHV216 resistant plasmid, each grown in competition with a susceptible *A. baylyi* BD413 strain with the opposite fluorescent marker. “wastewater”: spiked and unspiked samples of either WWTP influent and effluent were inoculated with a 1:1 ratio of both strains and incubated overnight at 37 °C before plating was performed. CFUs were counted after incubation of 48 h at 30 °C. “rebuilt concentrations”: the antibiotic concentrations measured in the wastewater or the spiked concentrations were rebuilt in 1:10-diluted Müller-Hinton broth (MHB) and then inoculated likewise. “control”: competition experiments were performed as controls in 1:10-diluted MHB without antibiotics. * = The CFU numbers in the wastewater samples with the plasmid carrying strains were very low (< 20), therefore, these results are less reliable. “n.g” = no growth on the agar plates. Standard deviations are shown in Figure S2 in the supplemental_1.

The concentrations of the selected target antibiotics measured in the WWTP influent and effluent, as well as the concentrations spiked into the WWTP influent and effluent samples (see Soufi et al. (2025) [65]), were added to 1:10-diluted MHB to test, whether the observed selection was caused by these antibiotics (rebuilt concentrations, Figure 1). The strain *A. baylyi* BD413 mCherry 777 and the plasmid-carrying *A. baylyi* BD413 strains reached a maximum of 11% of the counted colonies (Figure 1). *A. baylyi* BD413 mCherry 652 was not selected either, reaching around 40% of the colonies in the rebuilt influent and effluent, which is comparable to the results of the controls without antibiotics. Interestingly, when the same concentrations of antibiotics, which had been added to the spiked wastewater, were added to the 1:10-diluted MHB, no growth of any *A. baylyi* BD413 strain was observed (Figure 1). Table S1 (supplemental_1) compares the measured or spiked concentrations and the MIC and MSC values.

### 3.3 Antibiotic concentrations measured in the soil extracts

The concentrations of the six target antibiotics were measured in all soil extracts (colloidal and truly dissolved) via LCMSMS (Acquity UPLC H-Class Plus - Xevo TQ-S micro, Waters, USA). The maximum concentrations of both extract types depending on the incubation time and the spiking level (across all soil types or the wastewater quality) are listed in Table 4. Except for SMX, measured concentrations in the colloidal phases were always higher than the concentrations in the truly dissolved phase. Heatmaps of the antibiotic concentrations in the different extracts are shown in the supplemental (supplemental_1, Figure S1). Additionally, the measured concentrations [ng/L] of all extracts are provided in the supplemental materials (supplemental_2). A comparison between the maximum antibiotic concentrations from the extracts and the MSC values of our *A. baylyi* BD413 system is available in the supplemental materials (supplemental_1, Table S2). The measured concentrations were well below the MSCs of the strain pair.

**Table 4:** Maximum concentrations [µg/L] of the six target antibiotics measured in the colloidal and truly dissolved extracts via UPLC-MS/MS. *: One measured AZI concentration was excluded in this table as it exceeded all other concentrations of the antibiotic by at least factor 10, which was attributed to a measurement error. The measured concentrations of all extracts are provided in the supplemental (supplemental_2).

| Maximum concentrations [µg/L] |  |  |  |  |  |  |  |  |
| --- | --- | --- | --- | --- | --- | --- | --- | --- |
|  | 4 days |  |  |  | 4 weeks |  |  |  |
|  | colloidal |  | truly dissolved |  | colloidal |  | truly dissolved |  |
| antibiotic | unspiked | spiked | unspiked | spiked | unspiked | spiked | unspiked | spiked |
| SMX | 0.14 | 1.35 | 1.98 | 15.06 | 0.14 | 1.13 | 0.72 | 13.17 |
| TRM | 0.42 | 2.98 | 0.03 | 0.35 | 0.27 | 1.22 | 0.02 | 0.08 |
| ERY | 0.03 | 0.13 | 0.01 | 0.02 | 0.06 | 0.07 | 0.01 | 0.02 |
| CLI | 0.02 | 0.21 | 0.01 | 0.11 | 0.01 | 0.11 | 0.03 | 0.03 |
| CIP | < 0.01 | 0.01 | < 0.01 | < 0.01 | 0.01 | 0.01 | < 0.01 | < 0.01 |
| AZI | 0.02 | 0.04 | < 0.01 | 0.01* | 0.02 | 0.03 | < 0.01 | < 0.01 |

### 3.4 Colloidal soil extracts exert strong selective effects

At the end of the incubation experiment, (see 2.1) the soil samples were mixed with distilled water and then used to produce two extracts. The colloidal extracts contained the soil colloids with the potentially bound pharmaceuticals, while the truly dissolved extracts contained only substances that were completely dissolved in the water (see 2.2). To test the selective effects of these extracts on ARB, competition experiments were performed using the same resistant and susceptible *A. baylyi* BD413 strains as used for the wastewater. The results are shown in Figure 2.

**Figure 2:**
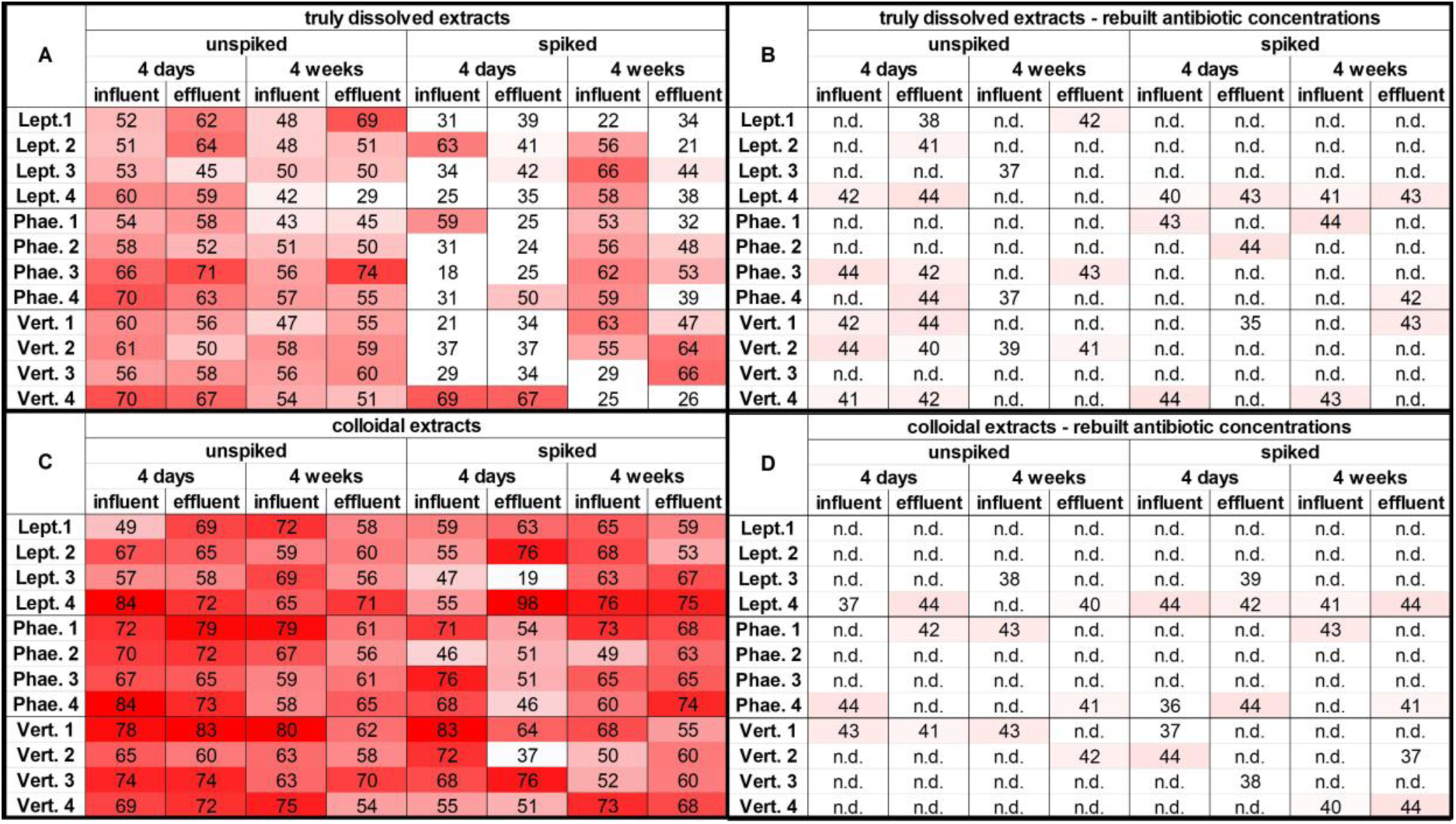
Heatmaps showing the percentage of growth (CFUs) of *A. baylyi* BD413 mCherry 652 grown in competition with *A. baylyi* BD413 GFP. Soil extracts (A, C) were inoculated with a 1:1 ratio of both strains and incubated overnight at 37 °C before plating was performed. CFUs were counted after incubation of 48 h at 30 °C. The antibiotic concentrations measured of selected extracts were rebuilt in 1:10-diluted Müller-Hinton broth and then inoculated likewise (B, D). “n.d.” = not determined. Standard deviations are shown in Figure S3 in the supplemental_1.

The resistant strain, *A. baylyi* BD413 mCherry 652 accounted for at least 51% of the counted CFUs in 58% of all truly dissolved extracts (Figure 3). While the amount of total CFUs was similar in the colloidal and the truly dissolved extracts, the selection of the resistant strain was stronger in the colloidal extracts than in the truly dissolved extracts. In the colloidal extracts, *A. baylyi* BD413 mCherry 652 accounted for 51% or more of the total CFUs in 92% of the extracts and outcompeted the susceptible *A. baylyi* BD413 GFP strain with at least 65% of the total CFUs in 53% of the extracts. This higher percentage was reached in 60% of the unspiked and in 46% of the spiked samples.

**Figure 3:**
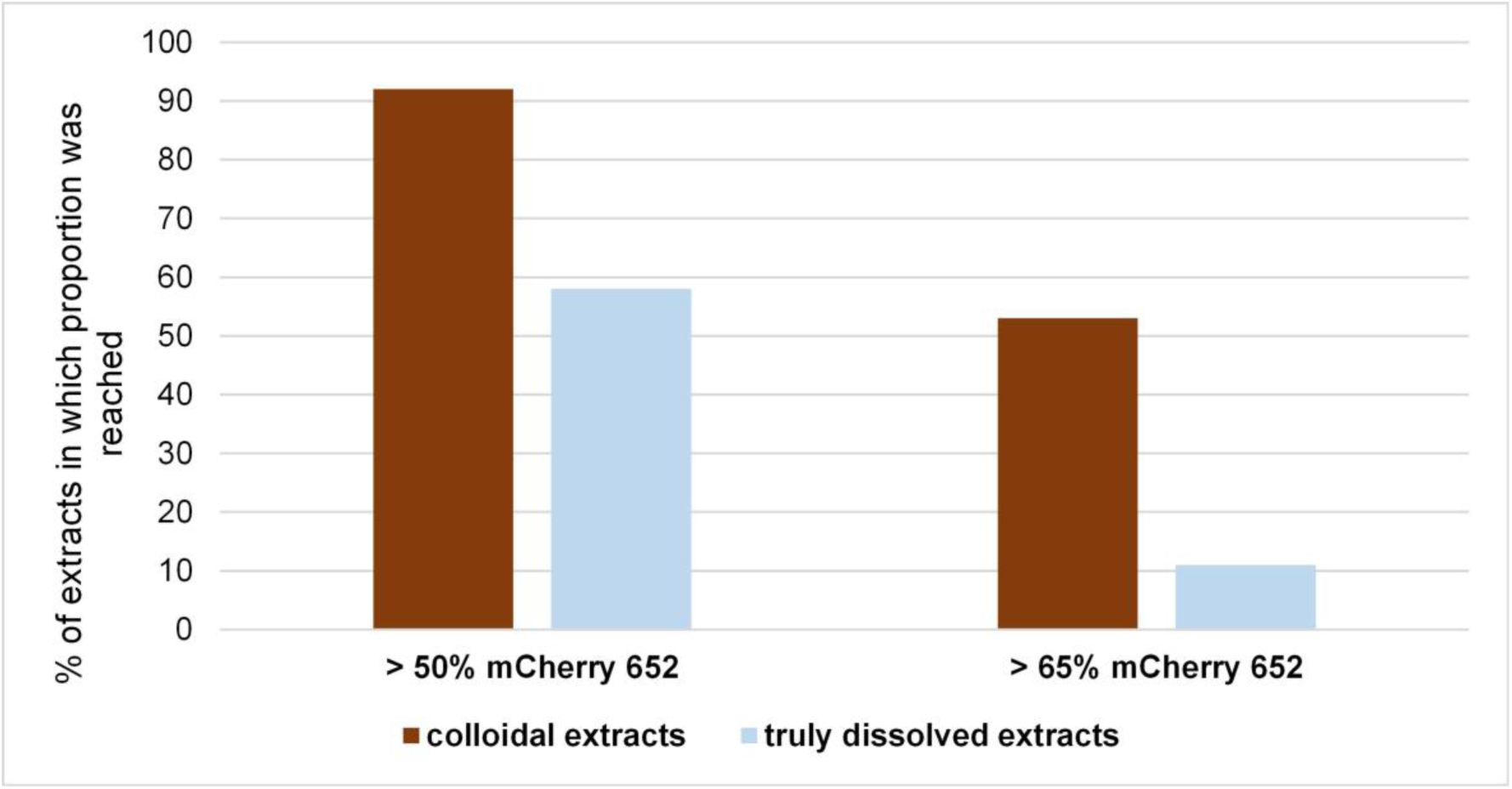
Proportional growth [%] of *A. baylyi* BD413 mCherry 652 in colloidal and truly dissolved extracts from soil-wastewater microcosms. Soil extracts were inoculated with a 1:1 ratio of the resistant *A. baylyi* BD413 mCherry 652 and the susceptible *A. baylyi* BD413 GFP and incubated overnight at 37 °C before plating was performed. CFUs were counted after incubation of 48 h at 30 °C.

The spiking level was the only factor with a significant effect (p < 0.0005) on the selection of the resistant strains in the colloidal and the truly dissolved extracts. Interestingly, the selection of the resistant strain was stronger in the unspiked extracts than in the spiked samples (colloidal and truly dissolved). The selection of the resistant strain tended to be stronger in extracts of soils irrigated with WWTP influent, reaching 63% compared to 44% in extracts of soils irrigated with effluent, but the difference was not statistically significant (p = 0.06). The other factors (soil type and incubation time) did also not show significant selection effects (p > 0.4). The highest percentage of *A. baylyi* BD413 mCherry 652 with 98% of all extracts (truly dissolved and colloidal) was reached in the sample from Leptosol 4, irrigated with spiked WWTP effluent after four days of incubation (Figure 2-C).

In both the truly dissolved and colloidal extracts, *A. baylyi* BD413 mCherry 777 and the strains carrying the pHHV216 plasmid did not outcompete the susceptible strain, reaching a maximum of 23% of the counted CFUs. These data are shown in the supplemental material (supplemental_1, Figure S4 and S5).

The concentrations of the selected antibiotics measured in the extracts were rebuilt in 1:10-diluted MHB for selected extracts. Competition experiments were conducted with *A. baylyi* BD413 mCherry 652 in competition with *A. baylyi* BD413 GFP to determine whether the observed selection was caused by these antibiotics. *A. baylyi* BD413 mCherry 652 reached around 40% of the CFUs in all rebuilt extracts (Figure 2- B & D), which is the same result as from the control experiments without added antibiotic. This indicates the presence of further selective substances in the extracts.

### 3.5 Non-targeted analysis reveals further antimicrobial substances

Of the six colloidal extracts we chose for a non-targeted analysis, three showed only weak selection for the resistant *A. baylyi* BD413 mCherry 652. The samples Leptosol 1 (unspiked, influent, 4 days), Phaeozem 2 (unspiked, effluent, 4 weeks), and Vertisol 4 (unspiked, effluent, 4 weeks) yielded in 48%, 56%, and 52% of the resistant strain, respectively. The other three extracts Leptosol 4 (unspiked, influent, 4 days), Phaeozem 4 (unspiked, effluent, 4 days), and Vertisol 1 (unspiked, effluent, 4 days) selected strongly for *A. baylyi* BD413 mCherry 652 resulting in 84%, 79%, and 83% of the resistant strain, respectively, in the competition experiments. In total 57 different substances were tentatively identified by the non-targeted screening (supplemental_3). Several antiparasitic drugs (albendazole, mebendazole, and thiabendazole), antifungal agents (azoxystrobin, carbendazim, and fluconazole), and insecticides (clothianidin, flubendiamide, imidacloprid, and DEET), as well as a broad spectrum of human medicinal drugs (e.g., antidepressants and heart medication), were detected. Most substances were abundant in both the weakly and strongly selecting extracts. However, while 28 of the 82 compounds were only present in the weakly selecting extracts, eight compounds were only detected in the strongly selecting extracts (namely flubendamide, PFOA, desethylatrazin, benzyl-phthalate, clothianidin, xanthosine, allopurinol and adenine). Interestingly, clarithromycin was found in all six extracts and nalidixic acid was found in all strong-selecting samples but only in half of the weak-selecting ones.

## 4. Discussion

### 4.1 WWTP influent and effluent select resistant *A. baylyi* BD413

ARBs are very frequently detected in wastewater and WWTPs [65,70–72]. They were also found in wastewater used for irrigation in the Mezquital Valley in Mexico [65]. Hospital wastewater contributes significantly to the introduction of ARBs and antibiotics into the overall wastewater system [63,71,73–75]. Kehl et al. (2022) [76] traced carbapenemase-producing *Enterobacteriaceae* from patient rooms in a maximum-care hospital to the WWTP’s effluent. While most studies demonstrate the abundance of ARBs to highlight the risk of dissemination into the environment, few investigate whether ARBs are further selected by the simultaneous presence of selecting antimicrobials. In contrast, we were able to observe the selection of our fluorescently labeled *A. baylyi* BD413 strains directly within the tested wastewater samples.

Of the strains tested, only *A. baylyi* BD413 mCherry 652 was able to outcompete the susceptible *A. baylyi* BD413 GFP strain in the unspiked influent and effluent samples (see Figure 1). Meanwhile, *A. baylyi* BD413 mCherry 777 and the pHHV216-carrying strains were not selected. Interestingly, there was little difference in the results between the influent and effluent samples. Soufi et al. (2025) [65] showed that the antibiotic concentrations and the bacterial abundance were only slightly reduced by the wastewater treatment of this WWTP. This may be due to the Atotonilco WWTP’s strategy of preserving higher nutrient levels in the treated water [42], or from the mixing of treated and untreated wastewater, as reported by Espira et al. (2024) [77]. Hence, we could confirm here that the missing capability of reducing antibiotic compounds in the WWTP will lead to an on-going selection of ARBs in the wastewater and hence also later in the receiving environment.

Efflux pumps are among the most common resistance mechanisms, as they export a broad spectrum of substrates from the cell, including several antibiotics, conferring resistance to multiple antibiotics simultaneously [78–80]. *A. baylyi* BD413 mCherry 652 has a deletion, and *A. baylyi* BD413 mCherry 777 has an amino acid exchange in the AdeN repressor of the AdeIJK multidrug efflux pump [63]. This efflux pump is present in all *Acinetobacter* strains and confers resistance to various antibiotics including ERY, TRM and fluoroquinolones [81]. All three antibiotics (ERY, TRM, and CIP, a fluoroquinolone) were present and additionally spiked into the influent and effluent wastewater samples [65], most likely promoting the selection of *A. baylyi* BD413 mCherry 652. The concentrations of AZI, SMX and CIP, which were spiked into the wastewater, exceeded the MSC values of *A. baylyi* BD413 mCherry 652 and *A. baylyi* BD413 mCherry 777, which might explain the high selection levels of both strains in the spiked wastewater samples. The spiked ERY concentration of 185 µg/L (see Table S1 from Soufi et al. (2025) [65]) was also within the range of the MSC of *A. baylyi* BD413 mCherry 652, highlighting the combined selection effect of antimicrobial mixtures [82].

Interestingly, reconstituting the concentrations of the six target antibiotics that were measured in the unspiked wastewater samples (influent and effluent), in 1:10-diluted MHB did not yield higher CFUs of *A. baylyi* BD413 mCherry 652 than the control without antibiotics (rebuilt concentrations, Figure 1). This indicates that the antibiotic concentrations determined in the unspiked wastewater, even combined, were insufficient to exert a selective effect. Reconstituting the antibiotic concentrations, that were spiked into the wastewater samples, in 1:10-diluted MHB inhibited the growth of all *A. baylyi* BD413 strains, because the spiked concentrations of CIP and AZI exceeded their respective MIC values (Table 2). Dorival-García et al. (2013) [83] demonstrated a high sorption of fluoroquinolones (including CIP) to suspended solids in sewage sludge. Similar results have been published for other antibiotics and pharmaceuticals in wastewater [62,84–88]. Therefore, the antibiotic concentrations, especially those of CIP and AZI spiked into the influent and effluent samples, were most likely partly adsorbed by the suspended solids and colloids in the wastewater. This enabled the growth of the added bacterial strains. For a direct comparison of the concentrations and the MICs and MSCs, see Table S1. However, the selection of the resistant strain in the spiked wastewater was stronger than in the unspiked samples. Thus, either the spiked antibiotics exceeded the sorption capabilities of the suspended solids in the wastewater, or they remained partly bioavailable despite the sorption.

### 4.2 Selection of resistant *A. baylyi* BD413 is stronger in colloidal extracts

Numerous studies have associated irrigation with untreated wastewater with an increase in ARBs and ARGs in the irrigated soil [27,31,89–92]. Some studies have found a positive correlation between the abundance of ARBs and ARGs and irrigation with reclaimed or treated wastewater [93,94], though other studies have not reported such a clear effect [95–98]. While most studies quantify the abundance of ARBs or ARGs in irrigated soils to indicate potential risk, few investigate whether co-occurring selective antimicrobials in irrigated soils cause the abundance, or if constant entry of new ARBs and ARGs due to wastewater is the cause. In contrast, our study set out to examine the selective effects of different soil-wastewater extracts on our fluorescently labeled *A. baylyi* BD413 strains.

Due to their high specific surface areas, colloids are known to bind to some pharmaceuticals including antibiotics [59,60]. They can also enhance the mobility of bound contaminants in soil and aqueous environments [99,100]. Some authors state that pharmaceuticals or disinfectants sorbed to colloids have a reduced bioavailability and are therefore of less risk for the environment [59,101–105]. In contrast to this, the selection of the resistant *A. baylyi* BD413 mCherry 652 was stronger in the colloidal extracts than in the truly dissolved extract. The resistant mutant reached more than 51% of the total CFUs in 92% of the colloidal extracts and only in 58% of the truly dissolved extracts. This may be attributed to the fact that some pharmaceuticals that select resistant strains accumulate in the colloidal phase. Hence, and in contrast to previous findings, our results show that contaminants bound to soil colloids can still impact bacterial growth and select for ARBs. It might even be the case that colloids foster the effectiveness when simultaneously serving as carbon and nutrient hotspots for microorganisms. Therefore, contaminants such as antibiotics with strong sorption affinities should not be considered less harmful than pollutants that remain in the aqueous phase under environmental conditions. Disinfectants of the group of QAACs capable of co- and cross selection for ARBs should also not be overlooked [106,107]. Instead, the role of colloidal binding of antibiotics and disinfectants for ARB and ARG selection should be elucidated in future studies and should then be considered in risk assessment.

Wastewater treatment did not significantly impact the selection of the resistant *A. baylyi* BD413 mCherry 652 (Figure 2), though there was a trend in the colloidal extracts towards slightly stronger selection of the resistant strains in extracts from soil irrigated with influent (p = 0.06). Apparently, the specific design of the Atotonilco WWTP [42] with subsequent mixing of untreated influent with the WWTP’s treated effluent [77] was fairly ineffective in reducing the selection potentials.

Surprisingly, no significant effect of the incubation time on the selection of the resistant strain was detected. Antibiotics can transform or degrade in soil over time due to various abiotic or biotic processes, such as hydrolysis, photodegradation, and microbial degradation, which depend heavily on specific soil properties and abiotic factors (e.g., temperature and pH) [108]. Further, an increased sequestration of antibiotics over time can reduce the bioavailability and hence the effectiveness of respective substances [109,110]. Indeed, a reduction of the concentrations after four weeks of incubation compared to four days was visible for most of our six target antibiotics. Still, the incubation time of the soil-wastewater extracts did not significantly impact the selection of the resistant strain in the competition experiments (p > 0.4). This, together with other arguments listed above indicates that possibly other substances in the wastewater are additionally responsible for the observed selection of the resistant strain, such as QAACs present at high concentrations, having high affinity to solids and potentially colloids and of known persistence and enrichment in soils [44]. As the selection was stronger in the colloidal extracts, we assume that these substances were enriched in the colloidal fraction, which additionally provides contact areas for the microbes.

### 4.3 Unknown compounds in wastewater-soil extracts select resistant strains

Surprisingly, the level of spiking in the wastewater had a significant effect on the selection of the resistant strain in the competition experiments (p < 0.0005). *A. baylyi* BD413 mCherry 652 consistently had higher CFU percentages in the unspiked samples than in spiked ones (see Figure 2). For the spiked samples, the measured concentrations in the influent and effluent of the six target antibiotics were artificially increased by a factor of 500 [65]. However, the measured concentrations in the spiked extracts were only 2-20 times higher than in the unspiked extracts (depending on the antibiotic), indicating that most spiked substances were absorbed by suspended solids in wastewater or by soil components. Reconstituting the measured concentrations of the six target antibiotics from selected extracts (spiked and unspiked) in 1:10-diluted MHB did not result in the selection (> 50% of the CFUs) of the resistant *A. baylyi* BD413 mCherry 652. The total CFU percentages reached by *A. baylyi* BD413 mCherry 652 in the rebuilt concentrations were similar to the percentages observed in competition experiments in 1:10-diluted MHB without added antibiotics, i.e., about 40 %. Indeed, even the highest measured concentrations of the six target antibiotics were far below the determined MSC values for *A. baylyi* BD413 GFP and *A. baylyi* BD413 mCherry 652. Therefore, the observed selection of the resistant strain cannot be explained by the concentrations of the six target antibiotics but rather must be attributed to other contaminants.

Soufi et al. (2025) [65] found that the total bacterial abundance was higher in soils irrigated with the spiked wastewater. This might indicate that bacteria that are tolerant or resistant to antibiotics and disinfectants used the spiked pharmaceuticals as a nutrient source. Especially, the abundance of *Pseudomonas* was elevated in the spiked soils [65]. This genus is known to have high tolerance due to efflux pumps against not only antibiotics and QAACs, but also pesticides (e.g., fungicides and insecticides), which are commonly used in agriculture [111]. The genus *Pseudomonas* is also associated with degradation of a broad range of pharmaceuticals, including QAACs [112,113], antibiotics (e.g., SMX and CIP) [114,115] and pesticides [116–119]. Therefore, one possible explanation for the weaker selection in the spiked extracts is that the spiked antibiotics and disinfectants inhibited the growth of susceptible microorganisms, leading to stronger growth of native species that can degrade selecting pharmaceuticals in the soil such as *Pseudomonas*. Due to the higher abundance of these species, the degree of the total degradation of all contaminants might be higher in the spiked samples than in the unspiked, which leads to less selection of the resistant *A. baylyi* BD413 strain. Indeed, a decrease of the measured antibiotic concentrations in the soil extracts after four weeks of incubation was evident (see Table 2 and supplemental_2). However, the total concentrations in the spiked fractions were still higher than in the unspiked fractions. Both results, the selection of the resistant strain as well as the stronger selection in the unspiked samples point to the fact, that the selection must have been influenced by compounds that are present in the wastewater and soil extracts, mainly in the colloidal fraction.

### 4.4 Numerous additional substances with potential for selection detected

The composition of pollutants in wastewater can be heterogeneous and complex, depending on the sources of the water. High concentrations of antibiotic residues and metabolites are commonly found in hospital effluents [63,120–122] and in effluents from antibiotic production facilities [32,123]. The presence and concentrations of antibiotics and other pharmaceuticals may vary greatly depending on human consumption and administration to livestock [124–128]. A broad range of pharmaceuticals and antibiotics was also detected in wastewater used for irrigation in the Mezquital Valley [41,45,47,65]. Our non-targeted screening revealed the presence of 82 different substances in total in the six colloidal extracts that were analyzed (see supplemental_3).

Clarithromycin belongs to the macrolide antibiotics. Resistance to the macrolide ERY often results in resistance against all other macrolides as well as against lincosamides and streptogramin B antibiotics [129,130]. The simultaneously presence of clarithromycin with ERY and AZI as two other macrolide antibiotics, as well as of A-ERY, a metabolite of ERY [131], increases the risk of macrolide resistance. The antibiotic nalidixic acid was detected in all replicates of the strongly selecting extracts. Nalidixic acid, the first quinolone antibiotic [132], was primarily used to treat uncomplicated urinary tract infections [133]. Due to its comparably high MICs, low serum concentrations, side effects, and rapidly evolving resistances, the antibiotic was replaced by other fluoroquinolones, such as CIP [133–136]. However, cross-resistance against the entire fluoroquinolone class is common (see reviews by Pham et al. (2019) [135] and Redgrave et al. (2014) [137]). The mutant strain of this study *A. baylyi* BD413 mCherry 652 has a deletion of the AdeN regulator of the AdeIJK efflux pump. This efflux pump is known to confer resistance also to macrolides and fluoroquinolones [81,138]. Therefore, selective effects of other macrolide antibiotics or fluoroquinolones in the competition experiments conducted here are highly probable.

In addition, the non-targeted analysis also detected non-antibiotic drugs (NADs), such as antidepressants, heart medication, and painkillers. Carbamazepine was detected in all six selected extracts. Wang et al. (2019) [139] demonstrated that this antiepileptic drug promotes the dissemination of ARGs via HGT due to activation of the SOS response in *Escherichia coli*. Similarly, an increased uptake of free DNA in the presence of different NADs (e.g., ibuprofen or naproxen) was demonstrated for *A. baylyi* ADP1 [140]. Some of these NADs were previously detected in the wastewater canals of the Mezquital Valley by Lesser et al. (2018) [45]. Numerous other NADs have also been shown to influence antibiotic resistance, as reviewed by Murray et al. (2024) [141]. The authors of the review attributed this partly to elevated efflux pump expression, which leads to cross-resistance against antibiotics, as well as to the synergistic effects of antibiotics and NADs [141].

Additionally, to antibiotics and NADs, different pesticides were found in the extracts, which may have influenced the selection process in our competition experiments. Several of these compounds were only detected in the strongly selecting extracts. Unlike antibiotics and other drugs, pesticides are applied directly to fields or crops, so concentrations may be higher than those of drugs applied via wastewater irrigation and vary between different fields. Pesticides, such as fungicides, herbicides, and insecticides, can increase antibiotic resistance by promoting HGT, which leads to the spread of existing ARGs. Pesticides can also lead to an increased mutation frequency, resulting in *de novo* mutations that confer antibiotic resistance [141–143]. Carbendazim and azoxystrobin, fungicides that were detected in our extracts, increased the abundance of different ARGs in greenhouse soils in microcosm experiments [144]. Azoxystrobin was also shown to affect soil communities’ resistance to aminoglycoside antibiotics. Interestingly, the impact was stronger in loamy sand soils than in clay loam soils [145], highlighting the significant influence of soil composition on selection processes.

The non-targeted analysis of the selected extracts revealed numerous substances that may have influenced our competition experiments. Further experiments are needed to verify which substances had a strong selecting effect towards the resistant *A. baylyi* BD413 mCherry and to determine the exact concentrations of the detected substances in all extracts, as well as the respective MIC and MSC values.

Finally, also QAACs may have played a role in selection effects in the incubation experiment and extracted colloids but could not be measured due to limitations in amounts of colloids. They should be included in future studies because of their strong association to surfaces and documented persistence in soils [146].

## 5. Conclusion

Competitive interactions between susceptible and resistant *A. baylyi* BD413 strains provide an integrative assessment of how wastewater irrigation shapes selective pressures for antibiotic resistance in agricultural soils. Competition experiments resulted in the strong selection of the resistant *A. baylyi* BD413 mCherry 652 in the wastewater samples and colloidal extracts from wastewater-irrigated soil, irrespective of wastewater treatment. Hence, our results do not support the hypothesis that a significant solubilization of antibiotics that accumulated in the soils due to past during irrigation with untreated wastewater occurred upon irrigation with WWTP influent; i.e., the environmental pressure that leads to a selection of ARB was maintained. The effect of wastewater treatment, in turn, was low for the design of this wastewater treatment process, aiming to keep nutrient concentrations in the effluent as high as possible. Unexpectedly, the truly dissolved soil extracts produced a weaker selective effect across all tested soil types and incubation conditions. Therefore, we reject the hypothesis that antibiotics bound to soil colloids are less effective than antibiotics in the truly dissolved phase. In contrast, colloid-associated antibiotics and most likely other pharmaceuticals remained bioavailable to the tested bacteria and exerted a selective effect toward antibiotic resistance. Therefore, future research on wastewater and wastewater-irrigated soils should also include the colloidal fraction and should not consider pharmaceuticals bound to colloids as deactivated.

Throughout the experiments, our results revealed that explanations based solely on antibiotic concentrations were insufficient to account for the observed selection patterns. Measurable antibiotic concentrations were far below the MSCs values and did not lead to the selection of resistant strains in control experiments. This indicates that direct antibiotic selection is unlikely to be the primary driver of resistance evolution in these systems. Non-targeted screening confirmed the presence of diverse other contaminants, including additional antibiotics, pesticides, and NADs associated with the selection of ARBs and ARGs. The ARBs and ARGs in wastewater-irrigated environments are likely shaped not only by the antibiotics, but also by broader contaminant mixtures. Current environmental risk assessment frameworks, which largely focus on single-compound antibiotic thresholds, may thus substantially underestimate the complexity and strength of resistance selection in real agricultural systems.

## Supporting information

Supplemental_1: All other figures and tables

Supplemental_2: Antibiotic concentrations

Supplemental_3: Results untargeted screening

## Acknowledgements/Funding

This research was funded by the Deutsche Forschungsgemeinschaft (DFG, German Research Foundation) in the framework of the Research Unit FOR 5095: Pollutant – Antibiotic Resistance – Pathogen Interactions in a Changing Wastewater Irrigation System, project number 431531292. The Mexican partners acknowledge funding by UNAM-DGAPA-PAPIIT project number AG101524 and AG101221 “Respuesta de las interacciones entre contaminantes, nutrientes y microorganismos al tratamiento del agua en los suelos del Valle del Mezquital y su efecto en la dispersión de multidrogoresistencia”. We would also like to thank Kathia Lüneberg for organizing the wastewater and soil sampling campaign. We thank Alessio Anselm and Alexander White for their assistance in the laboratory. We also thank Leila Soufi, Sara Gallego, Dipen Pulami, Stefanie Glaeser, Elisabeth Grohmann, and Kornelia Smalla for helping set up the soil microcosms for the incubation experiment.

## CRediT authorship contribution statement

**Katharina Axtmann:** Conceptualization, Data curation, Formal analysis, Investigation, Methodology, Writing – original draft, Writing – review & editing. **Christina Paffenholz:** Methodology. **Alina Auerhammer:** Methodology. **Ana Karen Michel-Farias:** Methodology. **Benjamin Justus Heyde:** Project administration, Conceptualization. **Lena Marie Coppers** Data curation. **Melanie Braun:** Supervision, Conceptualization. **Arne Kappenberg:** Methodology. **Susanne Brüggen:** Methodology, Data curation. **Ines Mulder:** Project administration, Conceptualization. **Christina Siebe:** Project administration, Funding acquisition, Conceptualization. **Wulf Amelung:** Project administration, Funding acquisition. **Jan Siemens:** Project administration, Funding acquisition, Conceptualization. **Gabriele Bierbaum:** Writing – review & editing, Supervision, Project administration, Funding acquisition, Conceptualization.

## Data availability

Additional data will be made available on request.

## References

[1] A. Fleming, On the Antibacterial Action of Cultures of a Penicillium, with Special Reference to their Use in the Isolation of B. influenzæ, British journal of experimental pathology 10 (1929) 226–236.

[2] I.M. Gould, A.M. Bal, New antibiotic agents in the pipeline and how they can help overcome microbial resistance, Virulence 4 (2013) 185–191. 10.4161/viru.22507.

[3] B. Spellberg, D.N. Gilbert, The future of antibiotics and resistance: a tribute to a career of leadership by John Bartlett, Clinical infectious diseases an official publication of the Infectious Diseases Society of America 59 Suppl 2 (2014) S71–5. 10.1093/cid/ciu392.

[4] M. Naghavi, S.E. Vollset, K.S. Ikuta, L.R. Swetschinski, A.P. Gray, E.E. Wool, G. Robles Aguilar, T. Mestrovic, G. Smith, C. Han, R.L. Hsu, J. Chalek, D.T. Araki, E. Chung, C. Raggi, A. Gershberg Hayoon, N. Davis Weaver, P.A. Lindstedt, A.E. Smith, U. Altay, N.V. Bhattacharjee, K. Giannakis, F. Fell, B. McManigal, N. Ekapirat, J.A. Mendes, T. Runghien, O. Srimokla, A. Abdelkader, S. Abd-Elsalam, et al.C.J.L. Murray, Global burden of bacterial antimicrobial resistance 1990–2021: a systematic analysis with forecasts to 2050, The Lancet 404 (2024) 1199–1226. 10.1016/S0140-6736(24)01867-1.

[5] J.S. Mackenzie, M. Jeggo, The One Health Approach-Why Is It So Important?, Tropical medicine and infectious disease 4 (2019). 10.3390/tropicalmed4020088.

[6] P. Musicha, T. Morse, D. Cocker, L. Mugisha, C.P. Jewell, N.A. Feasey, Time to define One Health approaches to tackling antimicrobial resistance, Nature communications 15 (2024) 8782. 10.1038/s41467-024-53057-z.

[7] R. Rohwedder, T. Bergan, S.B. Thorsteinsson, H. Scholl, Transintestinal elimination of ciprofloxacin, Chemotherapy 36 (1990) 77–84. 10.1159/000238751.

[8] V.K. Gupta, G. Maier, L. Gasink, A. Ek, M. Fudeman, P. Srivastava, A. Talley, Absorption, Metabolism, and Excretion of 14C-Tebipenem Pivoxil Hydrobromide (TBP-PI-HBr) in Healthy Male Subjects, Antimicrobial agents and chemotherapy 67 (2023) e0150922. 10.1128/aac.01509-22.

[9] S. Miyazaki, T. Katsube, H. Shen, C. Tomek, Y. Narukawa, Metabolism, Excretion, and Pharmacokinetics of 14 C-Cefiderocol (S-649266), a Siderophore Cephalosporin, in Healthy Subjects Following Intravenous Administration, Journal of clinical pharmacology 59 (2019) 958–967. 10.1002/jcph.1386.

[10] M. Anwar, Q. Iqbal, F. Saleem, Improper disposal of unused antibiotics: an often overlooked driver of antimicrobial resistance, Expert review of anti-infective therapy 18 (2020) 697–699. 10.1080/14787210.2020.1754797.

[11] D.G.J. Larsson, C. de Pedro, N. Paxeus, Effluent from drug manufactures contains extremely high levels of pharmaceuticals, Journal of hazardous materials 148 (2007) 751–755. 10.1016/j.jhazmat.2007.07.008.

[12] M. Milaković, G. Vestergaard, J.J. González-Plaza, I. Petrić, A. Šimatović, I. Senta, S. Kublik, M. Schloter, K. Smalla, N. Udiković-Kolić, Pollution from azithromycin-manufacturing promotes macrolide-resistance gene propagation and induces spatial and seasonal bacterial community shifts in receiving river sediments, Environment international 123 (2019) 501–511. 10.1016/j.envint.2018.12.050.

[13] F. Ju, K. Beck, X. Yin, A. Maccagnan, C.S. McArdell, H.P. Singer, D.R. Johnson, T. Zhang, H. Bürgmann, Wastewater treatment plant resistomes are shaped by bacterial composition, genetic exchange, and upregulated expression in the effluent microbiomes, The ISME journal 13 (2019) 346–360. 10.1038/s41396-018-0277-8.

[14] T.U. Berendonk, C.M. Manaia, C. Merlin, D. Fatta-Kassinos, E. Cytryn, F. Walsh, H. Bürgmann, H. Sørum, M. Norström, M.-N. Pons, N. Kreuzinger, P. Huovinen, S. Stefani, T. Schwartz, V. Kisand, F. Baquero, J.L. Martinez, Tackling antibiotic resistance: the environmental framework, Nature reviews. Microbiology 13 (2015) 310–317. 10.1038/nrmicro3439.

[15] M.S. Kostich, A.L. Batt, J.M. Lazorchak, Concentrations of prioritized pharmaceuticals in effluents from 50 large wastewater treatment plants in the US and implications for risk estimation, Environmental pollution (Barking, Essex 1987) 184 (2014) 354–359. 10.1016/j.envpol.2013.09.013.

[16] B. Heyde, M. Braun, L. Soufi, K. Lüneberg, S. Gallego, W. Amelung, K. Axtmann, G. Bierbaum, S.P. Glaeser, E. Grohmann, R. Arredondo-Hernández, I. Mulder, D. Pulami, K. Smalla, C. Zarfl, C. Siebe, J. Siemens, Transition from irrigation with untreated wastewater to treated wastewater and associated benefits and risks, npj Clean Water 8 (2025). 10.1038/s41545-025-00438-6.

[17] O. Paltiel, G. Fedorova, G. Tadmor, G. Kleinstern, Y. Maor, B. Chefetz, Human Exposure to Wastewater-Derived Pharmaceuticals in Fresh Produce: A Randomized Controlled Trial Focusing on Carbamazepine, Environmental science & technology 50 (2016) 4476–4482. 10.1021/acs.est.5b06256.

[18] T.J. Downs, E. Cifuentes-García, I.M. Suffet, Risk screening for exposure to groundwater pollution in a wastewater irrigation district of the Mexico City region, Environmental health perspectives 107 (1999) 553–561. 10.1289/ehp.99107553.

[19] C.B. Chigor, I.-A.I. Ibangha, N.O. Nweze, V.C. Onuora, C.A. Ozochi, Y. Titilawo, M.C. Enebe, T.N. Chernikova, P.N. Golyshin, V.N. Chigor, Prevalence of integrons in multidrug-resistant Escherichia coli isolates from waters and vegetables in Nsukka and Enugu, Southeast Nigeria, Environmental science and pollution research international 29 (2022) 60945–60952. 10.1007/s11356-022-20254-6.

[20] P. Drechsel, M. Qadir, D. Galibourg, The WHO Guidelines for Safe Wastewater Use in Agriculture: A Review of Implementation Challenges and Possible Solutions in the Global South, Water 14 (2022) 864. 10.3390/w14060864.

[21] Q.-A. Ahmad, E. Moors, H. Biemans, N. Shaheen, I. Masih, M.Z. ur Rahman Hashmi, Climate-induced shifts in irrigation water demand and supply during sensitive crop growth phases in South Asia, Climatic Change 176 (2023). 10.1007/s10584-023-03629-7.

[22] A. Beltran-Peña, P. D’Odorico, Future Food Security in Africa Under Climate Change, Earth’s Future 10 (2022). 10.1029/2022EF002651.

[23] G.G. Haile, Q. Tang, K.W. Reda, B. Baniya, L. He, Y. Wang, S.H. Gebrechorkos, Projected impacts of climate change on global irrigation water withdrawals, Agricultural Water Management 305 (2024) 109144. 10.1016/j.agwat.2024.109144.

[24] E.L. Williams, J.T. Abatzoglou, Climate Change Increases Evaporative and Crop Irrigation Demand in North America, Earth’s Future 13 (2025). 10.1029/2025EF005931.

[25] P. Dalkmann, M. Broszat, C. Siebe, E. Willaschek, T. Sakinc, J. Huebner, W. Amelung, E. Grohmann, J. Siemens, Accumulation of pharmaceuticals, Enterococcus, and resistance genes in soils irrigated with wastewater for zero to 100 years in central Mexico, PloS one 7 (2012) e45397. 10.1371/journal.pone.0045397.

[26] J.C. Durán-Alvarez, E. Becerril-Bravo, V.S. Castro, B. Jiménez, R. Gibson, The analysis of a group of acidic pharmaceuticals, carbamazepine, and potential endocrine disrupting compounds in wastewater irrigated soils by gas chromatography-mass spectrometry, Talanta 78 (2009) 1159–1166. 10.1016/j.talanta.2009.01.035.

[27] K. Lüneberg, B. Prado, M. Broszat, P. Dalkmann, D. Díaz, J. Huebner, W. Amelung, Y. López-Vidal, J. Siemens, E. Grohmann, C. Siebe, Water flow paths are hotspots for the dissemination of antibiotic resistance in soil, Chemosphere 193 (2018) 1198–1206. 10.1016/j.chemosphere.2017.11.143.

[28] Y. Bigott, S. Gallego, N. Montemurro, M.-C. Breuil, S. Pérez, A. Michas, F. Martin-Laurent, P. Schröder, Fate and impact of wastewater-borne micropollutants in lettuce and the root-associated bacteria, The Science of the total environment 831 (2022) 154674. 10.1016/j.scitotenv.2022.154674.

[29] A. Christou, P. Karaolia, E. Hapeshi, C. Michael, D. Fatta-Kassinos, Long-term wastewater irrigation of vegetables in real agricultural systems: Concentration of pharmaceuticals in soil, uptake and bioaccumulation in tomato fruits and human health risk assessment, Water research 109 (2017) 24–34. 10.1016/j.watres.2016.11.033.

[30] F.O. Gudda, M.G. Waigi, E.S. Odinga, B. Yang, L. Carter, Y. Gao, Antibiotic-contaminated wastewater irrigated vegetables pose resistance selection risks to the gut microbiome, Environmental pollution (Barking, Essex 1987) 264 (2020) 114752. 10.1016/j.envpol.2020.114752.

[31] S. Jechalke, M. Broszat, F. Lang, C. Siebe, K. Smalla, E. Grohmann, Effects of 100 years wastewater irrigation on resistance genes, class 1 integrons and IncP-1 plasmids in Mexican soil, Frontiers in microbiology 6 (2015) 163. 10.3389/fmicb.2015.00163.

[32] J. Bengtsson-Palme, D.G.J. Larsson, Concentrations of antibiotics predicted to select for resistant bacteria: Proposed limits for environmental regulation, Environment international 86 (2016) 140–149. 10.1016/j.envint.2015.10.015.

[33] L. Serwecińska, Antimicrobials and Antibiotic-Resistant Bacteria: A Risk to the Environment and to Public Health, Water 12 (2020) 3313. 10.3390/w12123313.

[34] E. Gullberg, S. Cao, O.G. Berg, C. Ilbäck, L. Sandegren, D. Hughes, D.I. Andersson, Selection of resistant bacteria at very low antibiotic concentrations, PLoS pathogens 7 (2011) e1002158. 10.1371/journal.ppat.1002158.

[35] E. Gullberg, L.M. Albrecht, C. Karlsson, L. Sandegren, D.I. Andersson, Selection of a multidrug resistance plasmid by sublethal levels of antibiotics and heavy metals, mBio 5 (2014) e01918–14. 10.1128/mBio.01918-14.

[36] J. Song, C. Rensing, P.E. Holm, M. Virta, K.K. Brandt, Comparison of Metals and Tetracycline as Selective Agents for Development of Tetracycline Resistant Bacterial Communities in Agricultural Soil, Environmental science & technology 51 (2017) 3040–3047. 10.1021/acs.est.6b05342.

[37] C. Seiler, T.U. Berendonk, Heavy metal driven co-selection of antibiotic resistance in soil and water bodies impacted by agriculture and aquaculture, Frontiers in microbiology 3 (2012) 399. 10.3389/fmicb.2012.00399.

[38] J. Ye, C. Rensing, J. Su, Y.-G. Zhu, From chemical mixtures to antibiotic resistance, Journal of environmental sciences (China) 62 (2017) 138–144. 10.1016/j.jes.2017.09.003.

[39] A. Sánchez-González, B. González-Méndez, C. Siebe, Uso del agua residual urbana en la agricultura: los casos de México y China, in: Yolanda Trapaga Delfín (Coord). América Latina y el Caribe – China. Recursos Naturales y Medio Ambiente, Primera Edición. México: Unión de Universidades de América Latina y el Caribe; 2013, pp. 59–70.

[40] E.M. García-Salazar, El agua residual como generadora del espacio de la actividad agrícola en el Valle del Mezquital, Hidalgo, México, Estudios Sociales 29 (2019). 10.24836/es.v29i54.741.

[41] J.C. Durán-Álvarez, B. Prado, R. Zanella, M. Rodríguez, S. Díaz, Wastewater surveillance of pharmaceuticals during the COVID-19 pandemic in Mexico City and the Mezquital Valley: A comprehensive environmental risk assessment, The Science of the total environment 900 (2023) 165886. 10.1016/j.scitotenv.2023.165886.

[42] A.-L. Garduño-Jiménez, J.-C. Durán-Álvarez, C.A. Ortori, S. Abdelrazig, D.A. Barrett, R.L. Gomes, Delivering on sustainable development goals in wastewater reuse for agriculture: Initial prioritization of emerging pollutants in the Tula Valley, Mexico, Water research 238 (2023) 119903. 10.1016/j.watres.2023.119903.

[43] M.E. Gutiérrez-Ruiz, I. Sommer, C. Siebe, Effects of land application of waste water from Mexico City on soil fertility and heavy metal accumulation: a bibliographical review, Environ. Rev. 3 (1995) 318–330. 10.1139/a95-017.

[44] B.J. Heyde, A. Anders, C. Siebe, J. Siemens, I. Mulder, Quaternary alkylammonium disinfectant concentrations in soils rise exponentially after long-term wastewater irrigation, Environ. Res. Lett. 16 (2021) 64002. 10.1088/1748-9326/abf0cf.

[45] L.E. Lesser, A. Mora, C. Moreau, J. Mahlknecht, A. Hernández-Antonio, A.I. Ramírez, H. Barrios-Piña, Survey of 218 organic contaminants in groundwater derived from the world’s largest untreated wastewater irrigation system: Mezquital Valley, Mexico, Chemosphere 198 (2018) 510–521. 10.1016/j.chemosphere.2018.01.154.

[46] C. Siebe, W.R. Fischer, Effect of long-term irrigation with untreated sewage effluents on soil properties and heavy metal adsorption of leptosols and vertisols in Central Mexico, J Plant Nutrition & Soil 159 (1996) 357–364. 10.1002/jpln.1996.3581590408.

[47] J. Siemens, G. Huschek, C. Siebe, M. Kaupenjohann, Concentrations and mobility of human pharmaceuticals in the world’s largest wastewater irrigation system, Mexico City-Mezquital Valley, Water research 42 (2008) 2124–2134. 10.1016/j.watres.2007.11.019.

[48] M. Broszat, H. Nacke, R. Blasi, C. Siebe, J. Huebner, R. Daniel, E. Grohmann, Wastewater irrigation increases the abundance of potentially harmful gammaproteobacteria in soils in Mezquital Valley, Mexico, Applied and environmental microbiology 80 (2014) 5282–5291. 10.1128/AEM.01295-14.

[49] World Health Organization & UN-Habitat, Progress on safe treatment and use of wastewater: piloting the monitoring methodology and initial findings for SDG indicator 6.3.1, World Health Organization (2018).

[50] D.J. Rodriguez, H.A. Serrano, A. Delgado, D. Nolasco, G. Saltiel, From Waste to Resource: Shifting paradigms for smarter wastewater interventionsin Latin America and the Caribbean., World Bank, Washington, DC (2020).

[51] M. Carrillo, G.C. Braun, C. Siebe, W. Amelung, J. Siemens, Desorption of sulfamethoxazole and ciprofloxacin from long-term wastewater-irrigated soils of the Mezquital Valley as affected by water quality, J Soils Sediments 16 (2016) 966–975. 10.1007/s11368-015-1292-2.

[52] B.J. Heyde, K. Lüneberg, N. Hahn, M. Braun, J. Siemens, C. Siebe, Nutrients, Metals, and Carbon in Soils Irrigated With Treated Versus Untreated Wastewater, Z. Pflanzenernähr. Bodenk. (2025). 10.1002/jpln.70042.

[53] J.R. Lead, K.J. Wilkinson, Aquatic Colloids and Nanoparticles: Current Knowledge and Future Trends, Environmental Chemistry 3 (2006) 159–171. 10.1071/EN06025.

[54] N. Gottselig, W. Amelung, J.W. Kirchner, R. Bol, W. Eugster, S.J. Granger, C. Hernández-Crespo, F. Herrmann, J.J. Keizer, M. Korkiakoski, H. Laudon, I. Lehner, S. Löfgren, A. Lohila, C.J.A. Macleod, M. Mölder, C. Müller, P. Nasta, V. Nischwitz, E. Paul-Limoges, M.C. Pierret, K. Pilegaard, N. Romano, M.T. Sebastià, M. Stähli, M. Voltz, H. Vereecken, J. Siemens, E. Klumpp, Elemental Composition of Natural Nanoparticles and Fine Colloids in European Forest Stream Waters and Their Role as Phosphorus Carriers, Global Biogeochemical Cycles 31 (2017) 1592–1607. 10.1002/2017GB005657.

[55] A. Missong, R. Bol, V. Nischwitz, J. Krüger, F. Lang, J. Siemens, E. Klumpp, Phosphorus in water dispersible-colloids of forest soil profiles, Plant Soil 427 (2018) 71–86. 10.1007/s11104-017-3430-7.

[56] A. Hartland, Lead JR, V.I. Slaveykova, D. O’Carroll, E. Valsami-Jones, The environmental significance of natural nanoparticles, 8th ed., 2013.

[57] T.M. Tsao, Y.M. Chen, M.K. Wang, Origin, separation and identification of environmental nanoparticles: a review, Journal of environmental monitoring JEM 13 (2011) 1156–1163. 10.1039/c1em10013k.

[58] H.K. Shon, S. Vigneswaran, S.A. Snyder, Effluent Organic Matter (EfOM) in Wastewater: Constituents, Effects, and Treatment, Critical Reviews in Environmental Science and Technology 36 (2006) 327–374. 10.1080/10643380600580011.

[59] S. Bagnis, M.F. Fitzsimons, J. Snape, A. Tappin, S. Comber, Processes of distribution of pharmaceuticals in surface freshwaters: implications for risk assessment, Environ Chem Lett 16 (2018) 1193–1216. 10.1007/s10311-018-0742-7.

[60] S.J. Klaine, P.J.J. Alvarez, G.E. Batley, T.F. Fernandes, R.D. Handy, D.Y. Lyon, S. Mahendra, M.J. McLaughlin, J.R. Lead, Nanomaterials in the environment: behavior, fate, bioavailability, and effects, Environmental toxicology and chemistry 27 (2008) 1825–1851. 10.1897/08-090.1.

[61] K. Yu, C. Sun, B. Zhang, M. Hassan, Y. He, Size-dependent adsorption of antibiotics onto nanoparticles in a field-scale wastewater treatment plant, Environmental pollution (Barking, Essex 1987) 248 (2019) 1079–1087. 10.1016/j.envpol.2019.02.090.

[62] K. Maskaoui, J.L. Zhou, Colloids as a sink for certain pharmaceuticals in the aquatic environment, Environmental science and pollution research international 17 (2010) 898–907. 10.1007/s11356-009-0279-1.

[63] D. Schuster, K. Axtmann, N. Holstein, C. Felder, A. Voigt, H. Färber, P. Ciorba, C. Szekat, A. Schallenberg, M. Böckmann, C. Zarfl, C. Neidhöfer, K. Smalla, M. Exner, G. Bierbaum, Antibiotic concentrations in raw hospital wastewater surpass minimal selective and minimum inhibitory concentrations of resistant Acinetobacter baylyi strains, Environmental microbiology 24 (2022) 5721–5733. 10.1111/1462-2920.16206.

[64] IUSS Working Group WRB, World Reference Base for Soil Resources. International soil classification system for naming soils and creating legends for soil maps. 4th edition., International Union of Soil Sciences (IUSS), Vienna, Austria (2022).

[65] L. Soufi, I.D. Kampouris, K. Lüneberg, B.J. Heyde, D. Pulami, S.P. Glaeser, C. Siebe, J. Siemens, K. Smalla, E. Grohmann, S. Gallego, Wastewater-borne pollutants influenced antibiotic resistance genes and mobile genetic elements in the soil without affecting the bacterial community composition in a changing wastewater irrigation system, Journal of hazardous materials 494 (2025) 138680. 10.1016/j.jhazmat.2025.138680.

[66] M. Braun, L. Juraschek, N. Siebers, J. Kruse, Natural Colloids in Soil: Effect of Storage and Extraction Conditions on Colloid Amount and Composition, Z. Pflanzenernähr. Bodenk. 188 (2025) 813–821. 10.1002/jpln.202400341.

[67] A. Kappenberg, L.M. Juraschek, Development of a LC-MS/MS Method for the Simultaneous Determination of the Mycotoxins Deoxynivalenol (DON) and Zearalenone (ZEA) in Soil Matrix, Toxins 13 (2021). 10.3390/toxins13070470.

[68] H. Heuer, C. Kopmann, C.T.T. Binh, E.M. Top, K. Smalla, Spreading antibiotic resistance through spread manure: characteristics of a novel plasmid type with low %G+C content, Environmental microbiology 11 (2009) 937–949. 10.1111/j.1462-2920.2008.01819.x.

[69] S. Brüggen, O.J. Schmitz, A New Concept for Regulatory Water Monitoring Via High-Performance Liquid Chromatography Coupled to High-Resolution Mass Spectrometry, J. Anal. Test. 2 (2018) 342–351. 10.1007/s41664-018-0081-5.

[70] T. Jäger, N. Hembach, C. Elpers, A. Wieland, J. Alexander, C. Hiller, G. Krauter, T. Schwartz, Reduction of Antibiotic Resistant Bacteria During Conventional and Advanced Wastewater Treatment, and the Disseminated Loads Released to the Environment, Frontiers in microbiology 9 (2018) 2599. 10.3389/fmicb.2018.02599.

[71] C. Bréchet, J. Plantin, M. Sauget, M. Thouverez, D. Talon, P. Cholley, C. Guyeux, D. Hocquet, X. Bertrand, Wastewater treatment plants release large amounts of extended-spectrum β-lactamase-producing Escherichia coli into the environment, Clinical infectious diseases an official publication of the Infectious Diseases Society of America 58 (2014) 1658–1665. 10.1093/cid/ciu190.

[72] M. Oliveira, P. Truchado, R. Cordero-García, M.I. Gil, M.A. Soler, A. Rancaño, F. García, A. Álvarez-Ordóñez, A. Allende, Surveillance on ESBL-Escherichia coli and Indicator ARG in Wastewater and Reclaimed Water of Four Regions of Spain: Impact of Different Disinfection Treatments, Antibiotics (Basel, Switzerland) 12 (2023). 10.3390/antibiotics12020400.

[73] L. Drieux, S. Haenn, L. Moulin, V. Jarlier, Quantitative evaluation of extended-spectrum β-lactamase-producing Escherichia coli strains in the wastewater of a French teaching hospital and relation to patient strain, Antimicrobial resistance and infection control 5 (2016) 9. 10.1186/s13756-016-0108-5.

[74] K. Endalamaw, S. Tadesse, Z. Asmare, D. Kebede, M. Erkihun, B. Abera, Antimicrobial resistance profile of bacteria from hospital wastewater at two specialized hospitals in Bahir Dar city, Ethiopia, BMC microbiology 24 (2024) 525. 10.1186/s12866-024-03693-8.

[75] A. Laffite, P.I. Kilunga, J.M. Kayembe, N. Devarajan, C.K. Mulaji, G. Giuliani, V.I. Slaveykova, J. Poté, Hospital Effluents Are One of Several Sources of Metal, Antibiotic Resistance Genes, and Bacterial Markers Disseminated in Sub-Saharan Urban Rivers, Frontiers in microbiology 7 (2016) 1128. 10.3389/fmicb.2016.01128.

[76] K. Kehl, A. Schallenberg, C. Szekat, C. Albert, E. Sib, M. Exner, N. Zacharias, C. Schreiber, M. Parčina, G. Bierbaum, Dissemination of carbapenem resistant bacteria from hospital wastewater into the environment, The Science of the total environment 806 (2022) 151339. 10.1016/j.scitotenv.2021.151339.

[77] L.M. Espira, J.D. Contreras, E.E. Felix-Arellano, C. Siebe, M. Mazari-Hiriart, H. Riojas-Rodríguez, J.N.S. Eisenberg, A comparative analysis of regional infection risk due to wastewater recontamination in the Mezquital Valley, Mexico, The Science of the total environment 919 (2024) 170615. 10.1016/j.scitotenv.2024.170615.

[78] A. Gaurav, P. Bakht, M. Saini, S. Pandey, R. Pathania, Role of bacterial efflux pumps in antibiotic resistance, virulence, and strategies to discover novel efflux pump inhibitors, Microbiology (Reading, England) 169 (2023). 10.1099/mic.0.001333.

[79] V. Thakur, A. Uniyal, V. Tiwari, A comprehensive review on pharmacology of efflux pumps and their inhibitors in antibiotic resistance, European journal of pharmacology 903 (2021) 174151. 10.1016/j.ejphar.2021.174151.

[80] M.A. Webber, L.J.V. Piddock, The importance of efflux pumps in bacterial antibiotic resistance, The Journal of antimicrobial chemotherapy 51 (2003) 9–11. 10.1093/jac/dkg050.

[81] L. Damier-Piolle, S. Magnet, S. Brémont, T. Lambert, P. Courvalin, AdeIJK, a resistance-nodulation-cell division pump effluxing multiple antibiotics in Acinetobacter baumannii, Antimicrobial agents and chemotherapy 52 (2007) 557–562. 10.1128/AAC.00732-07.

[82] M. Böckmann, K. Axtmann, G. Bierbaum, C. Zarfl, Pulsed antibiotic release into the environment may foster the spread of antimicrobial resistance, FEMS microbiology ecology 102 (2025). 10.1093/femsec/fiaf128.

[83] N. Dorival-García, A. Zafra-Gómez, A. Navalón, J. González, J.L. Vílchez, Removal of quinolone antibiotics from wastewaters by sorption and biological degradation in laboratory-scale membrane bioreactors, The Science of the total environment 442 (2013) 317–328. 10.1016/j.scitotenv.2012.10.026.

[84] Y. Aminot, L. Fuster, P. Pardon, K. Le Menach, H. Budzinski, Suspended solids moderate the degradation and sorption of waste water-derived pharmaceuticals in estuarine waters, The Science of the total environment 612 (2018) 39–48. 10.1016/j.scitotenv.2017.08.162.

[85] Q. Cui, J. Liu, Y. Tang, Y. Ma, G. Lin, R. Wang, W. Zhang, Q. Zuo, X. Zhao, F. Wu, Study of the adsorption behavior of tetracycline onto suspended sediments in the Yellow River, China: Insights into the transportation and mechanism, The Science of the total environment 889 (2023) 164242. 10.1016/j.scitotenv.2023.164242.

[86] G.G. Hacıosmanoğlu, M. Arenas, C. Mejías, J. Martín, J.L. Santos, I. Aparicio, E. Alonso, Adsorption of Fluoroquinolone Antibiotics from Water and Wastewater by Colemanite, International journal of environmental research and public health 20 (2023). 10.3390/ijerph20032646.

[87] R.D. Holbrook, N.G. Love, J.T. Novak, Sorption of 17beta-estradiol and 17alpha-ethinylestradiol by colloidal organic carbon derived from biological wastewater treatment systems, Environmental science & technology 38 (2004) 3322–3329. 10.1021/es035122g.

[88] H. Tu, J. Gao, Di Su, Y. Wang, J. Gao, Y. Wang, H. Li, Q. Liao, Y. Zheng, Differential Adsorption Behaviors of Light and Heavy SPM Fractions on Three Antibiotics: Implications for Lacustrine Antibiotic Migration, Water 17 (2025) 1859. 10.3390/w17131859.

[89] A. Malik, A. Aleem, Incidence of metal and antibiotic resistance in Pseudomonas spp. from the river water, agricultural soil irrigated with wastewater and groundwater, Environmental monitoring and assessment 178 (2011) 293–308. 10.1007/s10661-010-1690-2.

[90] O.A. Palacios, C.A. Contreras, L.N. Muñoz-Castellanos, M.O. González-Rangel, H. Rubio-Arias, A. Palacios-Espinosa, G.V. Nevárez-Moorillón, Monitoring of indicator and multidrug resistant bacteria in agricultural soils under different irrigation patterns, Agricultural Water Management 184 (2017) 19–27. 10.1016/j.agwat.2017.01.001.

[91] M. Pan, L.M. Chu, Occurrence of antibiotics and antibiotic resistance genes in soils from wastewater irrigation areas in the Pearl River Delta region, southern China, The Science of the total environment 624 (2018) 145–152. 10.1016/j.scitotenv.2017.12.008.

[92] S. Shafiani, A. Malik, Tolerance of pesticides and antibiotic resistance in bacteria isolated from wastewater-irrigated soil, World Journal of Microbiology and Biotechnology 19 (2003) 897–901. 10.1023/B:WIBI.0000007290.94694.4f.

[93] I.D. Kampouris, U. Klümper, S. Agrawal, L. Orschler, D. Cacace, S. Kunze, T.U. Berendonk, Treated wastewater irrigation promotes the spread of antibiotic resistance into subsoil pore-water, Environment international 146 (2021) 106190. 10.1016/j.envint.2020.106190.

[94] O.A. Palacios, F.J. La Zavala-Díaz de Serna, M.d.L. Ballinas-Casarrubias, M.S. Espino-Valdés, G.V. Nevárez-Moorillón, Microbiological Impact of the Use of Reclaimed Wastewater in Recreational Parks, International journal of environmental research and public health 14 (2017). 10.3390/ijerph14091009.

[95] F. Cerqueira, V. Matamoros, J. Bayona, G. Elsinga, L.M. Hornstra, B. Piña, Distribution of antibiotic resistance genes in soils and crops. A field study in legume plants (Vicia faba L.) grown under different watering regimes, Environmental research 170 (2019) 16–25. 10.1016/j.envres.2018.12.007.

[96] F. Cerqueira, V. Matamoros, J. Bayona, B. Piña, Antibiotic resistance genes distribution in microbiomes from the soil-plant-fruit continuum in commercial Lycopersicon esculentum fields under different agricultural practices, The Science of the total environment 652 (2019) 660–670. 10.1016/j.scitotenv.2018.10.268.

[97] Y. Negreanu, Z. Pasternak, E. Jurkevitch, E. Cytryn, Impact of treated wastewater irrigation on antibiotic resistance in agricultural soils, Environmental science & technology 46 (2012) 4800–4808. 10.1021/es204665b-Abstract.

[98] E. Troiano, L. Beneduce, A. Gross, Z. Ronen, Antibiotic-Resistant Bacteria in Greywater and Greywater-Irrigated Soils, Frontiers in microbiology 9 (2018) 2666. 10.3389/fmicb.2018.02666.

[99] J. Xu, R. Marsac, D. Costa, W. Cheng, F. Wu, J.-F. Boily, K. Hanna, Co-Binding of Pharmaceutical Compounds at Mineral Surfaces: Molecular Investigations of Dimer Formation at Goethite/Water Interfaces, Environmental science & technology 51 (2017) 8343–8349. 10.1021/acs.est.7b02835.

[100] S.B. Roy, D.A. Dzombak, Chemical Factors Influencing Colloid-Facilitated Transport of Contaminants in Porous Media, Environ. Sci. Technol. 31 (1997) 656–664. 10.1021/es9600643.

[101] B.J. Heyde, S.P. Glaeser, L. Bisping, K. Kirchberg, R. Ellinghaus, J. Siemens, I. Mulder, Smectite clay minerals reduce the acute toxicity of quaternary alkylammonium compounds towards potentially pathogenic bacterial taxa present in manure and soil, Scientific reports 10 (2020) 15397. 10.1038/s41598-020-71720-5.

[102] R. Cela-Dablanca, A. Barreiro, L. Rodríguez-López, P. Pérez-Rodríguez, M. Arias-Estévez, M.J. Fernández-Sanjurjo, E. Álvarez-Rodríguez, A. Núñez-Delgado, Azithromycin Adsorption onto Different Soils, Processes 10 (2022) 2565. 10.3390/pr10122565.

[103] K. Kümmerer, Antibiotics in the aquatic environment--a review--part I, Chemosphere 75 (2009) 417–434. 10.1016/j.chemosphere.2008.11.086.

[104] K. Lin, J. Gan, Sorption and degradation of wastewater-associated non-steroidal anti-inflammatory drugs and antibiotics in soils, Chemosphere 83 (2011) 240–246. 10.1016/j.chemosphere.2010.12.083.

[105] E. Topp, J. Renaud, M. Sumarah, L. Sabourin, Reduced persistence of the macrolide antibiotics erythromycin, clarithromycin and azithromycin in agricultural soil following several years of exposure in the field, The Science of the total environment 562 (2016) 136–144. 10.1016/j.scitotenv.2016.03.210.

[106] E. Walters, K. McClellan, R.U. Halden, Occurrence and loss over three years of 72 pharmaceuticals and personal care products from biosolids-soil mixtures in outdoor mesocosms, Water research 44 (2010) 6011–6020. 10.1016/j.watres.2010.07.051.

[107] S. Lennartz, J. Koschorreck, B. Göckener, K. Weinfurtner, A. Frohböse-Körner, J. Siemens, S. Balachandran, S.P. Glaeser, I. Mulder, Downstream effects of the pandemic? Spatiotemporal trends of quaternary ammonium compounds in suspended particulate matter of German rivers, Journal of hazardous materials 480 (2024) 136237. 10.1016/j.jhazmat.2024.136237.

[108] M. Cycoń, A. Mrozik, Z. Piotrowska-Seget, Antibiotics in the Soil Environment-Degradation and Their Impact on Microbial Activity and Diversity, Frontiers in microbiology 10 (2019) 338. 10.3389/fmicb.2019.00338.

[109] M. Förster, V. Laabs, M. Lamshöft, J. Groeneweg, S. Zühlke, M. Spiteller, M. Krauss, M. Kaupenjohann, W. Amelung, Sequestration of manure-applied sulfadiazine residues in soils, Environ. Sci. Technol. 43 (2009) 1824–1830. 10.1021/es8026538.

[110] I. Rosendahl, J. Siemens, J. Groeneweg, E. Linzbach, V. Laabs, C. Herrmann, H. Vereecken, W. Amelung, Dissipation and sequestration of the veterinary antibiotic sulfadiazine and its metabolites under field conditions, Environ. Sci. Technol. 45 (2011) 5216–5222. 10.1021/es200326t.

[111] N.M. Al-Enazi, M.S. AlTami, E. Alhomaidi, Unraveling the potential of pesticide-tolerant Pseudomonas sp. augmenting biological and physiological attributes of Vigna radiata (L.) under pesticide stress, RSC advances 12 (2022) 17765–17783. 10.1039/d2ra01570f.

[112] M. Tandukar, S. Oh, U. Tezel, K.T. Konstantinidis, S.G. Pavlostathis, Long-term exposure to benzalkonium chloride disinfectants results in change of microbial community structure and increased antimicrobial resistance, Environmental science & technology 47 (2013) 9730–9738. 10.1021/es401507k.

[113] U. Tezel, M. Tandukar, R.J. Martinez, P.A. Sobecky, S.G. Pavlostathis, Aerobic biotransformation of n-tetradecylbenzyldimethylammonium chloride by an enriched Pseudomonas spp. community, Environ. Sci. Technol. 46 (2012) 8714–8722. 10.1021/es300518c.

[114] B. Jiang, A. Li, Di Cui, R. Cai, F. Ma, Y. Wang, Biodegradation and metabolic pathway of sulfamethoxazole by Pseudomonas psychrophila HA-4, a newly isolated cold-adapted sulfamethoxazole-degrading bacterium, Applied microbiology and biotechnology 98 (2014) 4671–4681. 10.1007/s00253-013-5488-3.

[115] L. Li, J. Liu, J. Zeng, J. Li, Y. Liu, X. Sun, L. Xu, L. Li, Complete Degradation and Detoxification of Ciprofloxacin by a Micro-/Nanostructured Biogenic Mn Oxide Composite from a Highly Active Mn2+-Oxidizing Pseudomonas Strain, Nanomaterials (Basel, Switzerland) 11 (2021). 10.3390/nano11071660.

[116] P.K. Arora, A. Srivastava, V.P. Singh, Degradation of 4-chloro-3-nitrophenol via a novel intermediate, 4-chlororesorcinol by Pseudomonas sp. JHN, Scientific reports 4 (2014) 4475. 10.1038/srep04475.

[117] S. Jilani, M.A. Khan, Biodegradation of Cypermethrin by Pseudomonas in a batch activated sludge process, Int. J. Environ. Sci. Technol. 3 (2006) 371–380. 10.1007/BF03325946.

[118] S.E. Maloney, A. Maule, A.R. Smith, Microbial transformation of the pyrethroid insecticides: permethrin, deltamethrin, fastac, fenvalerate, and fluvalinate, Applied and environmental microbiology 54 (1988) 2874–2876. 10.1128/aem.54.11.2874-2876.1988.

[119] E. Topp, M.H. Akhtar, Identification and characterization of a pseudomonas strain capable of metabolizing phenoxybenzoates, Applied and environmental microbiology 57 (1991) 1294–1300. 10.1128/aem.57.5.1294-1300.1991.

[120] S. Aydin, M.E. Aydin, A. Ulvi, H. Kilic, Antibiotics in hospital effluents: occurrence, contribution to urban wastewater, removal in a wastewater treatment plant, and environmental risk assessment, Environmental science and pollution research international 26 (2019) 544–558. 10.1007/s11356-018-3563-0.

[121] M.D. Ekwanzala, R.F. Lehutso, T.K. Kasonga, J.B. Dewar, M.N.B. Momba, Environmental Dissemination of Selected Antibiotics from Hospital Wastewater to the Aquatic Environment, Antibiotics (Basel, Switzerland) 9 (2020). 10.3390/antibiotics9070431.

[122] T.Q. La Lien, N.Q. Hoa, N.T.K. Chuc, N.T.M. Thoa, H.D. Phuc, V. Diwan, N.T. Dat, A.J. Tamhankar, C.S. Lundborg, Antibiotics in Wastewater of a Rural and an Urban Hospital before and after Wastewater Treatment, and the Relationship with Antibiotic Use-A One Year Study from Vietnam, International journal of environmental research and public health 13 (2016). 10.3390/ijerph13060588.

[123] D.G.J. Larsson, Pollution from drug manufacturing: review and perspectives, Philosophical transactions of the Royal Society of London. Series B, Biological sciences 369 (2014). 10.1098/rstb.2013.0571.

[124] P. Gago-Ferrero, A.A. Bletsou, D.E. Damalas, R. Aalizadeh, N.A. Alygizakis, H.P. Singer, J. Hollender, N.S. Thomaidis, Wide-scope target screening of 2000 emerging contaminants in wastewater samples with UPLC-Q-ToF-HRMS/MS and smart evaluation of its performance through the validation of 195 selected representative analytes, Journal of hazardous materials 387 (2020) 121712. 10.1016/j.jhazmat.2019.121712.

[125] V. Hinnenkamp, P. Balsaa, T.C. Schmidt, Target, suspect and non-target screening analysis from wastewater treatment plant effluents to drinking water using collision cross section values as additional identification criterion, Analytical and bioanalytical chemistry 414 (2022) 425–438. 10.1007/s00216-021-03263-1.

[126] A. Kotwani, J. Joshi, D. Kaloni, Pharmaceutical effluent: a critical link in the interconnected ecosystem promoting antimicrobial resistance, Environmental science and pollution research international 28 (2021) 32111–32124. 10.1007/s11356-021-14178-w.

[127] J.A. Rivera-Jaimes, C. Postigo, R.M. Melgoza-Alemán, J. Aceña, D. Barceló, M. López de Alda, Study of pharmaceuticals in surface and wastewater from Cuernavaca, Morelos, Mexico: Occurrence and environmental risk assessment, The Science of the total environment 613–614 (2018) 1263–1274. 10.1016/j.scitotenv.2017.09.134.

[128] K. Wang, T. Zhuang, Z. Su, M. Chi, H. Wang, Antibiotic residues in wastewaters from sewage treatment plants and pharmaceutical industries: Occurrence, removal and environmental impacts, The Science of the total environment 788 (2021) 147811. 10.1016/j.scitotenv.2021.147811.

[129] B. Weisblum, Erythromycin resistance by ribosome modification, Antimicrobial agents and chemotherapy 39 (1995) 577–585. 10.1128/AAC.39.3.577.

[130] B. Weisblum, V. Demohn, Erythromycin-inducible resistance in Staphylococcus aureus: survey of antibiotic classes involved, Journal of bacteriology 98 (1969) 447–452. 10.1128/jb.98.2.447-452.1969.

[131] P. Kurath, P.H. Jones, R.S. Egan, T.J. Perun, Acid degradation of erythromycin A and erythromycin B, Experientia 27 (1971) 362. 10.1007/BF02137246.

[132] G.Y. Lesher, E.J. Froelich, M.D. Gruett, J.H. Bailey, R.P. Brundage, 1,8-Napththyridine derivatives. A new class of chemotherapeutic agents, Journal of medicinal and pharmaceutical chemistry 5 (1962) 1063–1065. 10.1021/jm01240a021.

[133] A.M. Emmerson, A.M. Jones, The quinolones: decades of development and use, The Journal of antimicrobial chemotherapy 51 Suppl 1 (2003) 13–20. 10.1093/jac/dkg208.

[134] M. Gellert, K. Mizuuchi, M.H. O’Dea, T. Itoh, J.I. Tomizawa, Nalidixic acid resistance: a second genetic character involved in DNA gyrase activity, Proceedings of the National Academy of Sciences of the United States of America 74 (1977) 4772–4776. 10.1073/pnas.74.11.4772.

[135] T.D.M. Pham, Z.M. Ziora, M.A.T. Blaskovich, Quinolone antibiotics, MedChemComm 10 (2019) 1719–1739. 10.1039/c9md00120d.

[136] R. Wise, Norfloxacin--a review of pharmacology and tissue penetration, The Journal of antimicrobial chemotherapy 13 Suppl B (1984) 59–64. 10.1093/jac/13.suppl_b.59.

[137] L.S. Redgrave, S.B. Sutton, M.A. Webber, L.J.V. Piddock, Fluoroquinolone resistance: mechanisms, impact on bacteria, and role in evolutionary success, Trends in microbiology 22 (2014) 438–445. 10.1016/j.tim.2014.04.007.

[138] E. Sugawara, H. Nikaido, Properties of AdeABC and AdeIJK efflux systems of Acinetobacter baumannii compared with those of the AcrAB-TolC system of Escherichia coli, Antimicrobial agents and chemotherapy 58 (2014) 7250–7257. 10.1128/AAC.03728-14.

[139] Y. Wang, J. Lu, L. Mao, J. Li, Z. Yuan, P.L. Bond, J. Guo, Antiepileptic drug carbamazepine promotes horizontal transfer of plasmid-borne multi-antibiotic resistance genes within and across bacterial genera, The ISME journal 13 (2019) 509–522. 10.1038/s41396-018-0275-x.

[140] Y. Wang, J. Lu, J. Engelstädter, S. Zhang, P. Ding, L. Mao, Z. Yuan, P.L. Bond, J. Guo, Non-antibiotic pharmaceuticals enhance the transmission of exogenous antibiotic resistance genes through bacterial transformation, The ISME journal 14 (2020) 2179–2196. 10.1038/s41396-020-0679-2.

[141] L.M. Murray, A. Hayes, J. Snape, B. Kasprzyk-Hordern, W.H. Gaze, A.K. Murray, Co-selection for antibiotic resistance by environmental contaminants, npj antimicrobials and resistance 2 (2024) 9. 10.1038/s44259-024-00026-7.

[142] Y. Xing, S. Wu, Y. Men, Exposure to Environmental Levels of Pesticides Stimulates and Diversifies Evolution in Escherichia coli toward Higher Antibiotic Resistance, Environ. Sci. Technol. 54 (2020) 8770–8778. 10.1021/acs.est.0c01155.

[143] D. Qiu, M. Ke, Q. Zhang, F. Zhang, T. Lu, L. Sun, H. Qian, Response of microbial antibiotic resistance to pesticides: An emerging health threat, The Science of the total environment 850 (2022) 158057. 10.1016/j.scitotenv.2022.158057.

[144] H. Zhang, S. Chen, Q. Zhang, Z. Long, Y. Yu, H. Fang, Fungicides enhanced the abundance of antibiotic resistance genes in greenhouse soil, Environmental pollution (Barking, Essex 1987) 259 (2020) 113877. 10.1016/j.envpol.2019.113877.

[145] M. Aleksova, A. Kenarova, S. Boteva, Azoxystrobin Impact on a Selection of Soil Bacterial Resistance to Amynoglicoside Antibiotics, “Prof. Marin Drinov” Publishing House of Bulgarian Academy of Sciences, 2019.

[146] K. Jansen, C. Mohr, K. Lügger, C. Heller, J. Siemens, I. Mulder, Widespread occurrence of quaternary alkylammonium disinfectants in soils of Hesse, Germany, The Science of the total environment 857 (2023) 159228. 10.1016/j.scitotenv.2022.159228.

