## Supplemental_1: All other figures and tables for "Wastewater and colloidal extracts of wastewater-irrigated soils select for resistant *Acinetobacter baylyi* beyond what measured antibiotic concentrations predict"

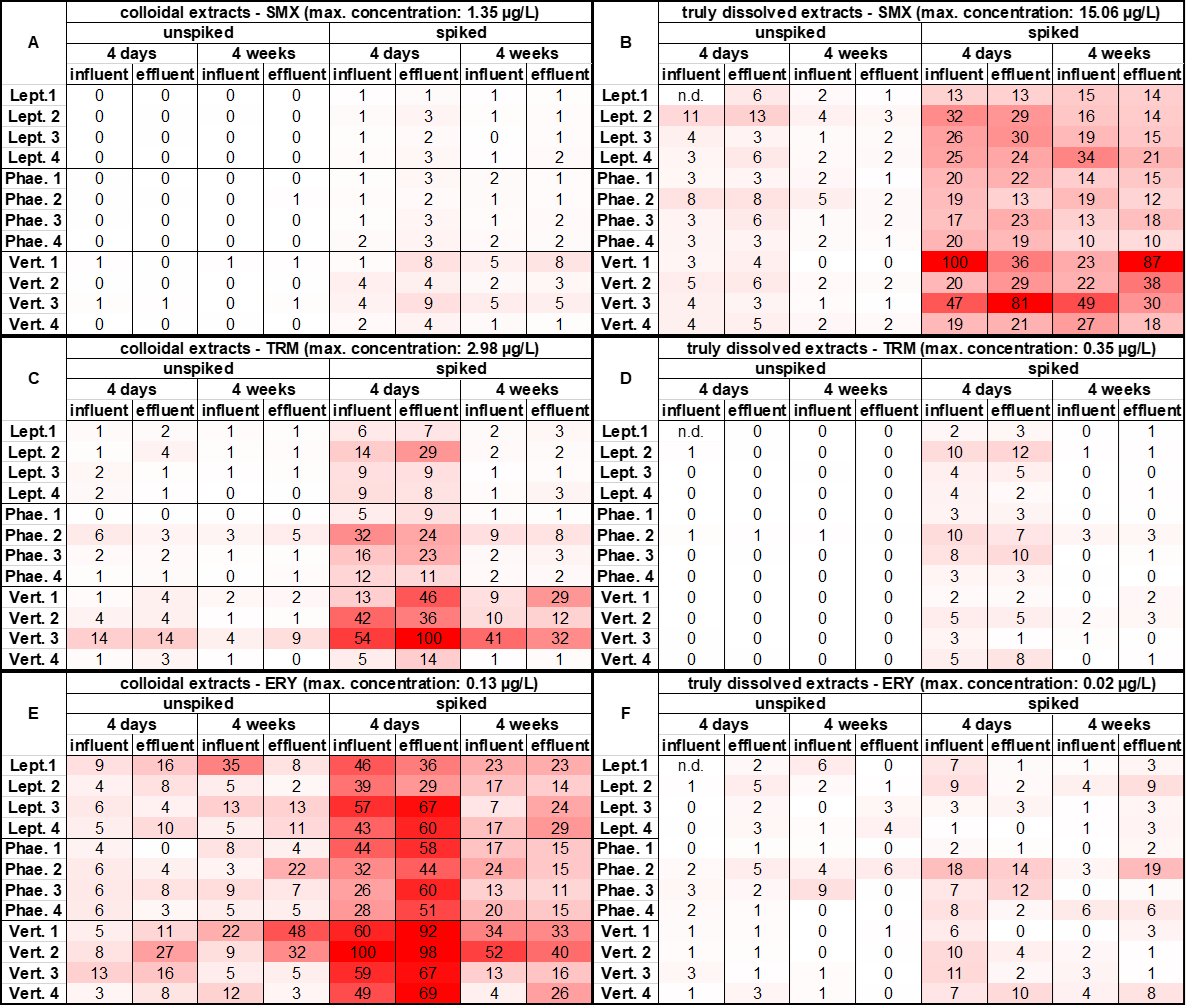

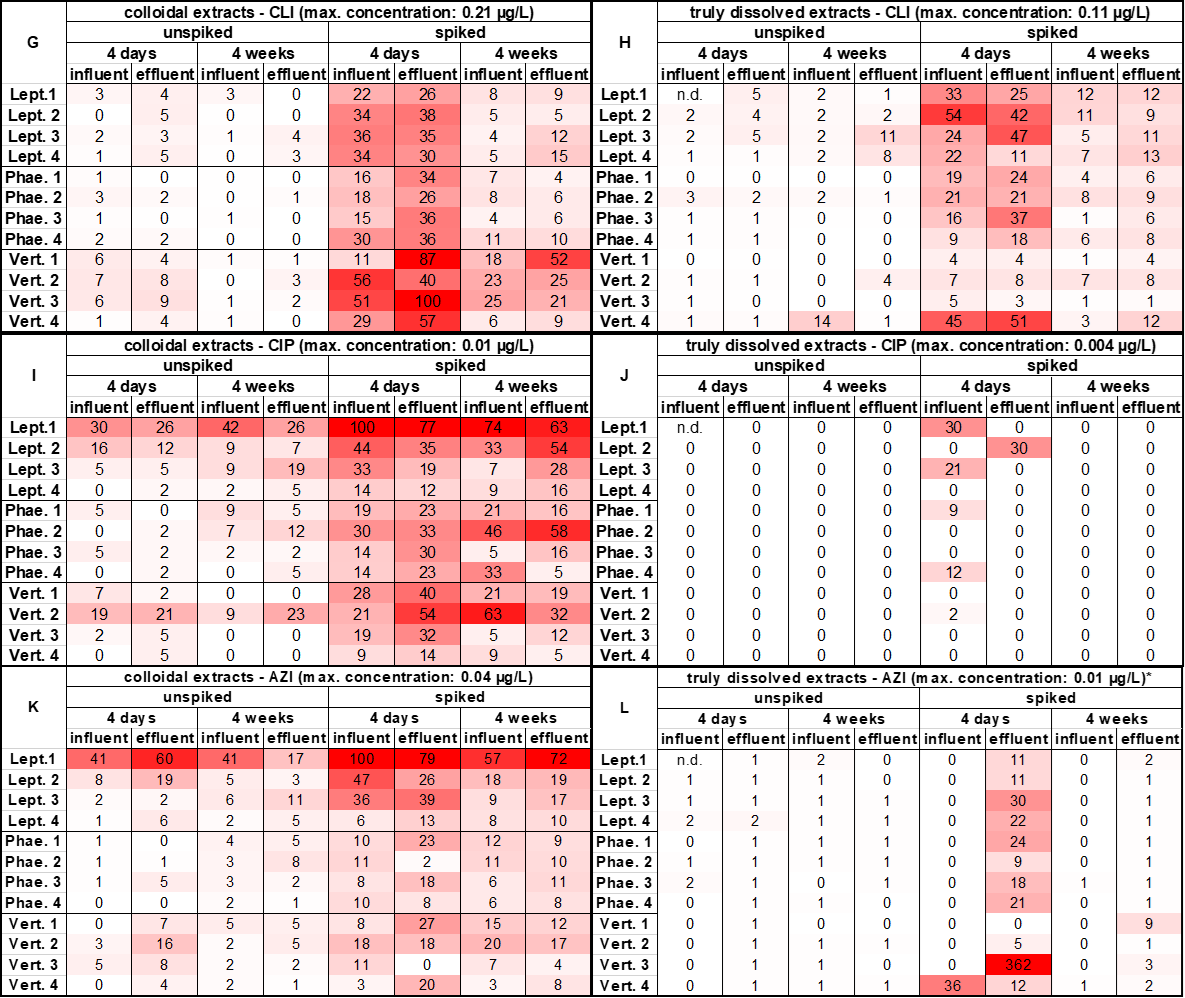

**Figure S1:** Heatmaps showing the relative concentrations [%] of the six target antibiotics compared to the respective highest value measured in the extracts, which is shown as “max. concentration” above the respective heatmap. A, C, E, G, H, I, K: colloidal extracts; B, D, F, H, J, L: truly dissolved extracts.

**Table S1:** Measured and spiked concentrations [µg/L] of the six target antibiotics from the WWTP influent and effluent in comparison with the MIC and MSC values of the susceptible and resistant A. baylyi BD413 strains. *: values determined and spiked by Soufi et al. (2025) [1]; **: as CLI was not detected, the spiked concentration was calculated on the base of concentrations published by Siemens et al. (2008) [2]; QL: quantification limit; DL: detection limit; ***: values determined by Schuster et al. (2022) [3]; “GFP” = A. baylyi BD413 GFP, “652” = A. baylyi BD413 mCherry 652, “777” = A. baylyi BD413 mCherry 777.

| Antibiotic [µg/L] | WWTP  influent* | WWTP  effluent* | Spiked  concentrations* | MIC GFP | MIC 652 | MIC 777 | MSC GFP & 652 | MSC GFP & 777 |
| --- | --- | --- | --- | --- | --- | --- | --- | --- |
| SMX | 3.19-3.99 | 2.82-3.78 | 1,995 | 3,000*** | 16,000*** | 128,000*** | 341.6 | - |
| TRM | 1.1-1.3 | 1.2-1.4 | 650 | 32,000 | >64,000 | >64,000 | 1799.7 | 2845.2 |
| ERY | 0.21 (<QL) -0.37 | 0.17-0.25 (<QL) | 185 | 2,000 | 16,000 | 8,000 | 255.5 | 559.7 |
| CLI | < DL | < DL | 60** | 8,000*** | 64,000*** | 64,000*** | 687.8*** | 1000.2*** |
| CIP | 2.35-3.39 | 1.94-2.27 | 1,695 | 20*** | 313*** | 313*** | 16.74*** | 51.5*** |
| AZI | 0-1.39 | 0-1.33 | 695 | 250 | 1,000 | 250 | 36.67 | - |

**Table S2:** Maximum concentrations [µg/L] of the six target antibiotics measured in the colloidal and truly dissolved extracts in comparison with the MSC values of the susceptible and resistant A. baylyi BD413 strains. *: One measured AZI concentration was excluded in this table as it exceeded all other concentrations by at least factor 10, which was attributed to a measurement error **: values determined by Schuster et al. (2022) [3]; “GFP” = A. baylyi BD413 GFP, “652” = A. baylyi BD413 mCherry 652, “777” = A. baylyi BD413 mCherry 777.

| Antibiotic [µg/L] | max. colloidal unspiked | max. colloidal spiked | max. truly dissolved unspiked | max. truly dissolved spiked | MSC GFP & 652 | MSC GFP & 777 |
| --- | --- | --- | --- | --- | --- | --- |
| SMX | 0.14 | 1.35 | 1.98 | 15.06 | 341.64 | - |
| TRM | 0.42 | 2.98 | 0.03 | 0.35 | 1799.7 | 2845.233 |
| ERY | 0.03 | 0.13 | 0.01 | 0.02 | 255.52 | 559.68 |
| CLI | 0.02 | 0.21 | 0.01 | 0.11 | 687.83** | 1000.22** |
| CIP | <0.01 | 0.01 | <0.01 | <0.01 | 16.74** | 51.51** |
| AZI | 0.02 | 0.04 | <0.01 | 0.01* | 36.67 | - |

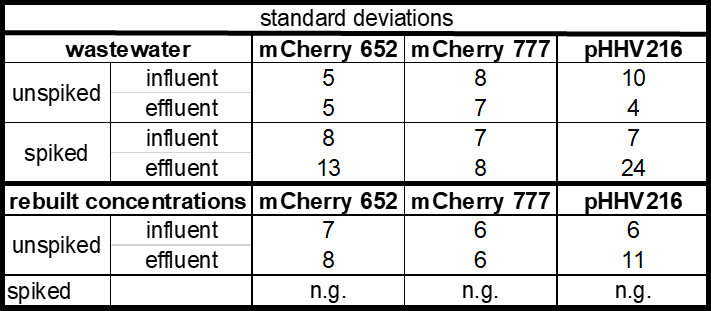

**Figure S2:** Standard deviations of the proportional growth [%] of A. baylyi BD413 mCherry 652, A. baylyi BD413 mCherry 777 or fluorescently labelled A. baylyi BD413 strains carrying the pHHV216 resistant plasmid, each grown in competition with a susceptible A. baylyi BD413 strain with the opposite fluorescent marker. Competition experiments were performed in wastewater (spiked and unspiked WWTP influent and effluent) and in 1:10-diluted MHB with reconstituted antibiotic concentrations from the wastewater samples. “n.g” = no growth on the agar plates.

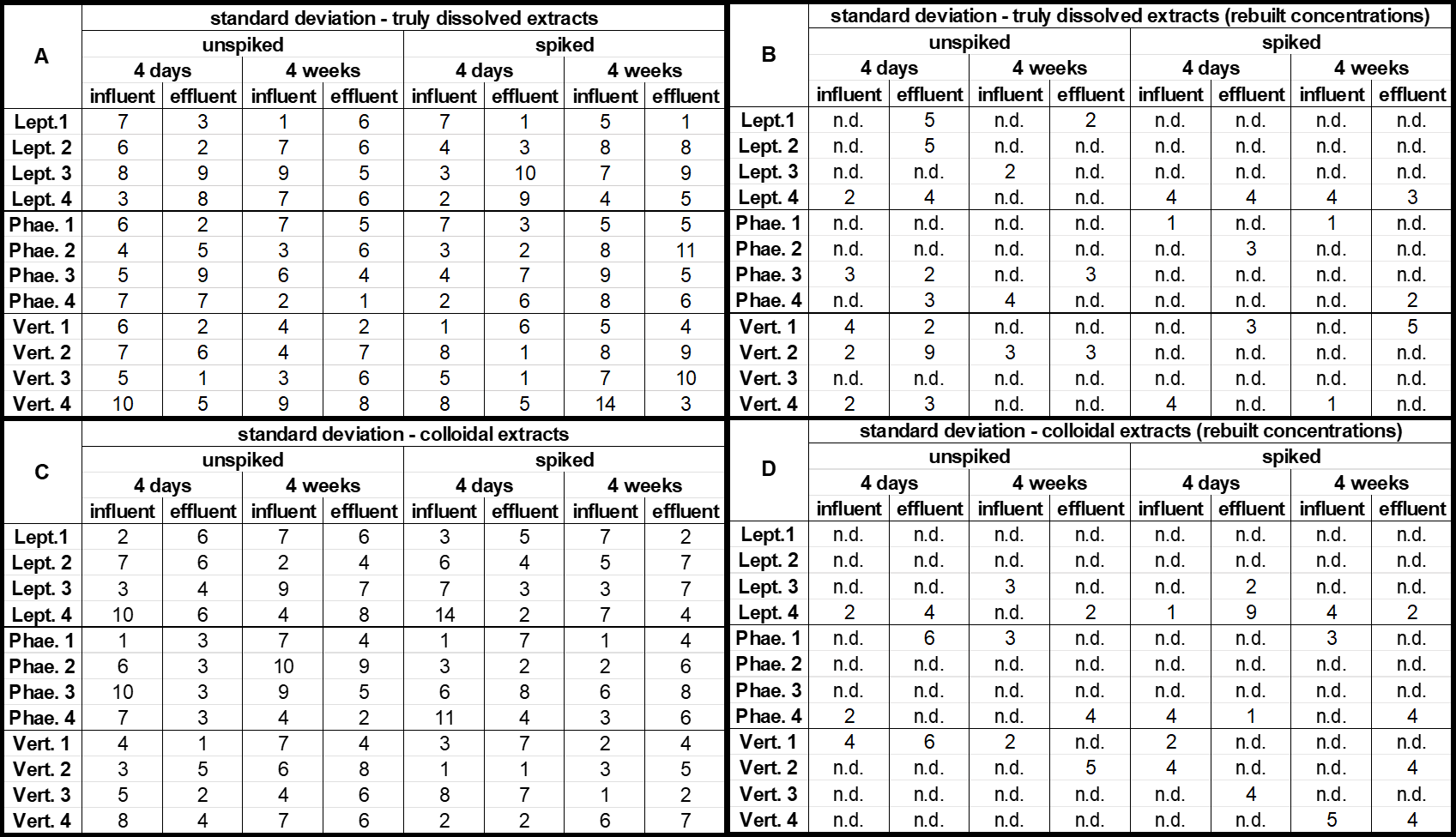

**Figure S3:** Standard deviations of the proportional growth [%] of A. baylyi BD413 mCherry 652 grown in competition with A. baylyi BD413 GFP shown in Figure 2 of the paper. Competition experiments were performed in soil extracts (A, C) and in 1:10-diluted MHB with reconstituted antibiotic concentrations from selected extracts (B, D). “n.d.” = not determined.

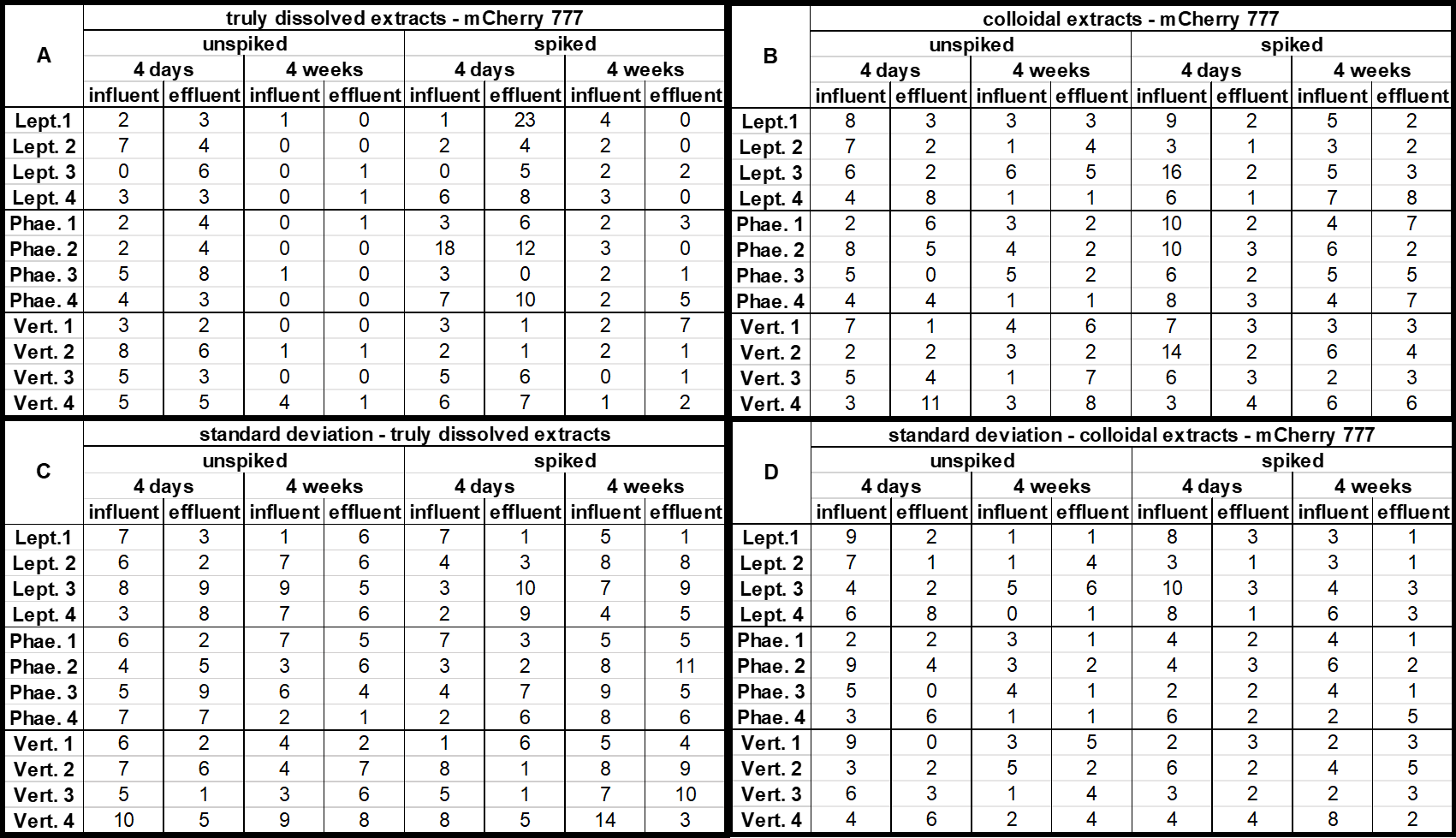

**Figure S4:** Percentage of A. baylyi BD413 mCherry 777 grown in competition with A. baylyi BD413 GFP on the total CFUs. Soil extracts (A, C) were inoculated with a 1:1 ratio of both strains and incubated overnight at 37 °C before plating was performed. CFUs were counted after incubation of 48 hours at 30 °C. B, D: Standard deviations of the respective competition experiments.

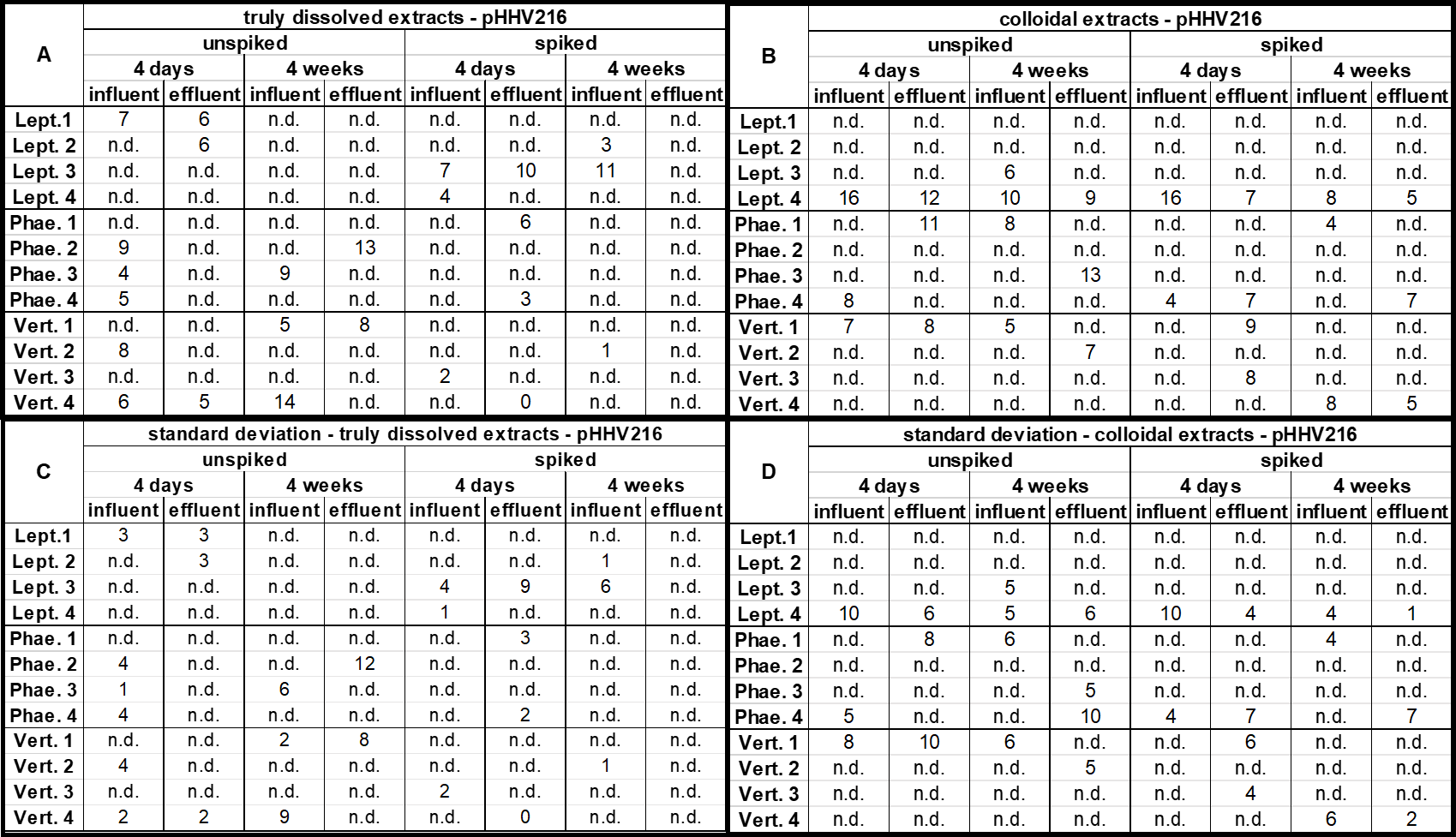

**Figure S5**: Percentage of A. baylyi BD413 carrying the pHHV216 resistance plasmid grown in competition with a susceptible A. baylyi BD413 strain with a different fluorescent marker on the total CFUs. Soil extracts (A, C) were inoculated with a 1:1 ratio of both strains and incubated overnight at 37 °C before plating was performed. CFUs were counted after incubation of 48 hours at 30 °C. B, D: Standard deviations of the respective competition experiments. “n.d.” = not determined.

**References**

[1]	L. Soufi, I.D. Kampouris, K. Lüneberg, B.J. Heyde, D. Pulami, S.P. Glaeser, C. Siebe, J. Siemens, K. Smalla, E. Grohmann, S. Gallego, Wastewater-borne pollutants influenced antibiotic resistance genes and mobile genetic elements in the soil without affecting the bacterial community composition in a changing wastewater irrigation system, Journal of hazardous materials 494 (2025) 138680. https://doi.org/10.1016/j.jhazmat.2025.138680.

[2]	J. Siemens, G. Huschek, C. Siebe, M. Kaupenjohann, Concentrations and mobility of human pharmaceuticals in the world's largest wastewater irrigation system, Mexico City-Mezquital Valley, Water research 42 (2008) 2124–2134. https://doi.org/10.1016/j.watres.2007.11.019.

[3]	D. Schuster, K. Axtmann, N. Holstein, C. Felder, A. Voigt, H. Färber, P. Ciorba, C. Szekat, A. Schallenberg, M. Böckmann, C. Zarfl, C. Neidhöfer, K. Smalla, M. Exner, G. Bierbaum, Antibiotic concentrations in raw hospital wastewater surpass minimal selective and minimum inhibitory concentrations of resistant Acinetobacter baylyi strains, Environmental microbiology 24 (2022) 5721–5733. https://doi.org/10.1111/1462-2920.16206.
